# Natural selection drives emergent genetic homogeneity in a century-scale experiment with barley

**DOI:** 10.1101/2023.09.22.557807

**Authors:** Jacob B Landis, Angelica M Guercio, Keely E Brown, Christopher J Fiscus, Peter L Morrell, Daniel Koenig

## Abstract

Direct observation is central to our understanding of the process of adaptation, but evolution is rarely documented in a large, multicellular organism for more than a few generations. Here, we observe genetic and phenotypic evolution across a century-scale competition experiment, barley composite cross II (CCII). CCII was founded in 1929 with tens of thousands of unique genotypes and has been adapted to local conditions in Davis, CA, USA for 58 generations. We find that natural selection has massively reduced genetic diversity leading to a single clonal lineage constituting most of the population by generation F50. Selection favored alleles originating from similar climates to that of Davis, and targeted genes regulating reproductive development, including some of the most well-characterized barley diversification loci, *Vrs1*, *HvCEN*, and *Ppd-H1*. We chronicle the dynamic evolution of reproductive timing in the population and uncover how parallel molecular pathways are targeted by stabilizing selection to optimize this trait. Our findings point to selection as the predominant force shaping genomic variation in one of the world’s oldest ongoing biological experiments.

**One-Sentence Summary:** Wholesale genetic restructuring of an experimental population is a consequence of rapid environmental adaptation.

---

The dispersal of cultivated plants is a classic example of rapid adaptive evolution <u>(*2*)</u>. Early farmers brought newly domesticated crops to wildly different environments, subjecting them to strong natural and artificial selective forces. These early crop varieties were phenotypically and genetically diverse, providing ample raw material to facilitate adaptation <u>(*3*)</u>. Ultimately, evolution resulted in locally adapted varieties that formed the backbone of early human civilization.

The history of the neolithic founder crop, barley (*Hordeum vulgare*), is a prototypical example of this process. Barley is a self-fertilizing, annual plant that was domesticated over 10,000 years ago and was dispersed from the Fertile Crescent to become a major source of nutrition for humans and livestock throughout Europe, Asia, and Northern Africa over just a few thousand generations <u>(*2*)</u>. Studies of collections of early cultivars have uncovered some of the population genetic history of the crop <u>(*4–7*)</u> and mapped loci that contributed to its spread <u>(*8–10*)</u>. However, precise estimates of the important parameters of genetic adaptation, such as the number and types of genes involved and the magnitude of selection at these loci, are limited without direct observation<u>(*11*)</u>.

We used a unique long-term competition experiment in barley to observe the process of local adaptation over decades. Composite cross II (CCII) is a multigenerational common garden experiment started in 1929 to adapt a genetically diverse population to the environmental conditions of Davis, CA, USA<u>(*1, 12*)</u>. Twenty-eight varieties were selected to represent the ecological, phenotypic, and geographical diversity of barley (Table S1) and were crossed in a half-diallel to generate 20,000 recombinant F2 progeny. Progeny from each cross were combined in equal proportion to found the experiment. Each year the population was propagated, 15,000- 20,000 seeds from CCII were sown, allowed to compete with minimal human intervention, and harvested in bulk. The harvested seeds were used in the following year to continue the experiment and live seeds of early generations were maintained by less frequent propagation of parallel lineages (Fig. 1, Fig S1).

**Fig. 1.**
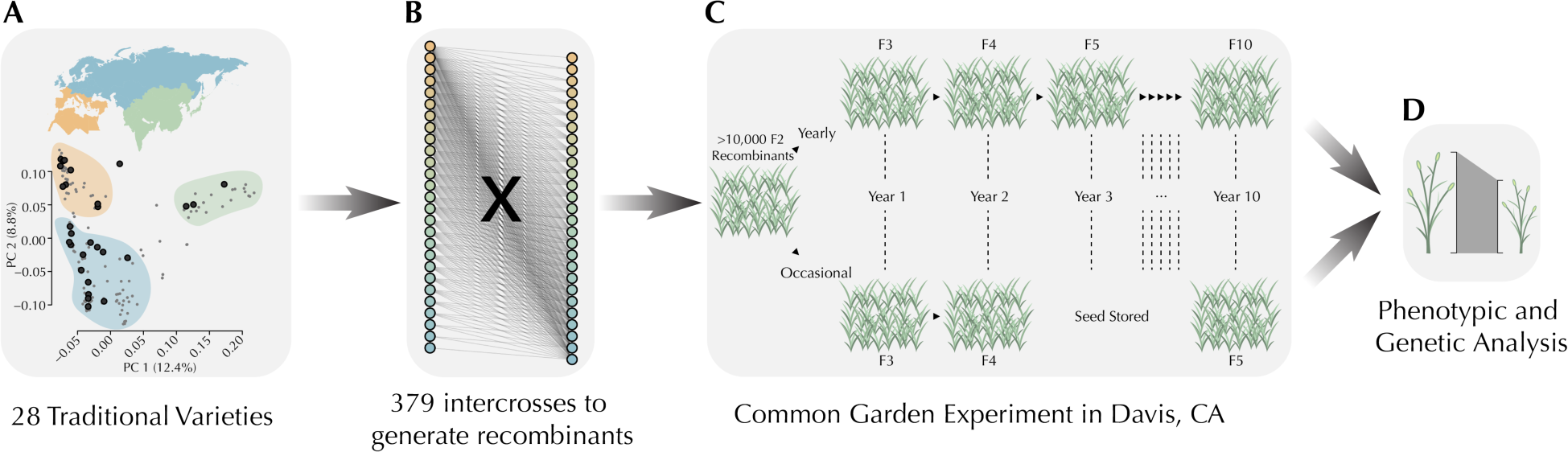
The design of composite cross II. (**A**) A principal component analysis of genetic diversity showing the distribution of the 28 CCII parents (dark grey) in a global panel of traditional varieties (light grey). (**B**) The crossing scheme of CCII where each of the 28 parents was crossed to each other parent in a single direction to generate 379 segregating families. (**C**) Illustration of the propagation of the CCII with less frequent grow outs of parallel lineages to retain live seed from earlier generations. (**D**) Seed reserved from previous grow outs can be used to compare changes over evolutionary time in phenotype and genotype.

Comparison of early and late generations of CCII has previously revealed substantial shifts in fitness-associated traits <u>(*12–15*)</u>. These include changes in the timing of reproduction, increases in plant biomass, and increases in seed size and number, indicating that natural selection was an important force shaping phenotypic diversity in CCII. Here, we leverage modern sequencing technologies to characterize the genetic underpinnings of adaptation over a half-century of evolution.

## The genetic diversity of CCII founders

We began by sequencing each of the genomes of the 28 CCII parents to a coverage depth of 8- 15x and identified 12,922,667 variants that segregated at the founding of the experiment.

Segregating variation in the founders included 64.8% of 1,316,845 SNPs segregating in a global survey of barley genetic diversity <u>(*10*)</u>, including 96.4% of common alleles (allele frequency > 0.1, Fig. S2), and estimated allele frequencies of genetic variants co-segregating in the parents and global datasets were well correlated (Spearman’s rho = 0.81, p < 2e-16, Fig. S3). Principal component analysis showed that CCII parents are well dispersed amongst global barley diversity along the first two PCs, representing 20.2% of the variance in the dataset (Fig. 1a, Fig S4). We conclude that the traditional cultivars selected to found CCII in the 1920’s reflect the distribution of common genetic variation in barley remarkably well.

## Rapid evolution in composite cross II

To understand how genetic diversity in CCII changed over time, we traced the evolutionary trajectory of each polymorphism from the founding population to generations F18, F28, and F58 (Fig S1) by sequencing pools of 1,000 individuals at each time point. Genetic diversity of CCII, as measured by mean expected heterozygosity (He) across all sites and the shape of the allele frequency spectrum, changed rapidly throughout the experiment (Fig. 2a, Fig S5). The rate of decrease of He was constant (-0.003 He / generation or 1.4% He^0^ / generation, p < 0.0012, R^2^ = 0.996) with a predicted genome-wide average of 0 by generation 71 at our sequencing depth. Allele frequency in the founding population was a strong predictor of eventual fixation genome-wide (Supplementary figure 5 and 6), with 78% of all alleles and 93% of rare alleles (MAF < 0.05) undetectable in F58. Near complete loss of genetic variation (He < 0.01) was initially restricted to a few regions of the genome (0.7% in F18), however by generations F50 and F58, 29.8% of the genome was near fixed at our sequencing depth. Average genome-wide FST was 0.18 between the founders and F58 (Fig. S7). Genome-wide FST was linearly associated with generational time increasing at a rate of 0.0027 ΔFST / generation (p < 0.003, R^2^ = 0.88, Fig. S7). The CCII lost genetic variation at a rate similar to that of neutral simulations conducted with population sizes two orders of magnitude smaller than the actual size of the experiment (Fig. 2a), much faster than expected due to genetic drift alone.

**Fig. 2.**
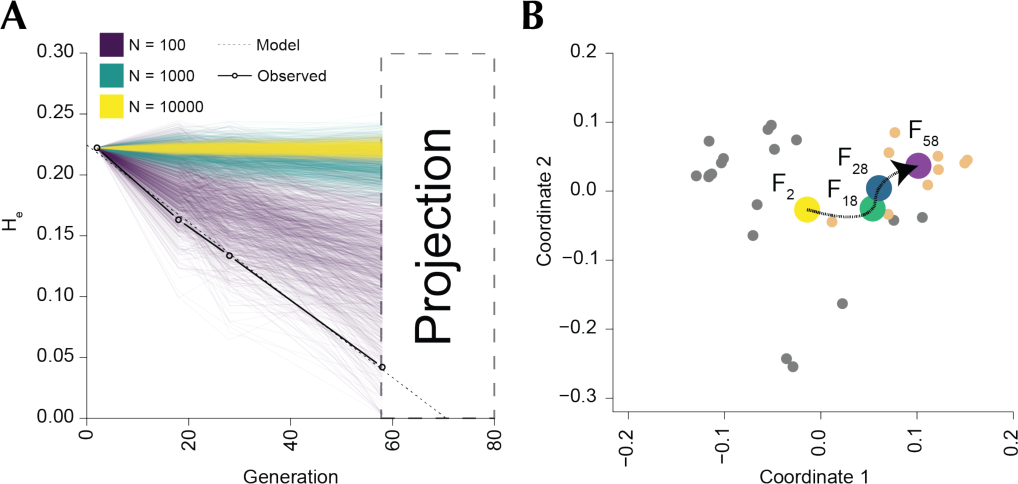
Rapid adaptive shift in the CCII. (**A**) Genome-wide loss of genetic diversity over time in the CCII compared to 1000 simulations of neutral evolution in the population using three different sizes. (**B**) Multi-dimensional scaling analysis showing the evolution of the CCII relative to the parents over time. Orange points are parents from Mediterranean climates in North Africa and grey are all other parents.

## A genome-wide signal of environmental adaptation

We considered two forms of selection that might be responsible for the unusually rapid genome-wide loss of diversity in CCII: selection against generally deleterious variants or positive selection for locally adapted alleles. While deleterious variation is not expected to show strong regional differences amongst barley varieties, locally adapted alleles would presumably be more common in parental lines that come from environments similar to Davis, CA.

To distinguish between these possibilities, we examined the evolution of CCII over time by calculating Nei’s genetic distance between genome-wide allele frequencies at each time point and the genotypes of each of the parental accessions (Fig. S8). We then used a multidimensional scaling procedure to determine the relationship of each CCII generation to the parents (Fig. 2b). CCII quickly evolved toward North African parents that grow in Mediterranean climates similar to that of Davis, CA. Examination of the genetic distances over time revealed this process to be dynamic, with the population most similar to the North African variety Arequipa collected in Peru, at time point F18, but in the later stages of the experiment, alleles derived from the variety Atlas were favored (Fig. S8). Atlas is a selection from the traditional variety brought by Spanish colonists to California in the late eighteenth century<u>(*16*)</u>.

The success of Mediterranean alleles was also evident for rare alleles that rose in frequency in the population. Minor alleles that increased in frequency were much more likely to be found in six-rowed North African parents (Fig. S8). Alleles identical to Atlas comprised 90.8% of fixed minor alleles (N = 779,883) and 82.2% of fixed private alleles (N = 10,739) in generation F58.

These results provide a picture of adaptation resembling historical experiments with barley that predate CCII<u>(*17*)</u>. These experiments competed barley varieties, rather than recombinants, at many sites throughout the United States. Maladapted varieties rapidly decreased in competition, allowing several varieties to increase in initial generations, but as time passed, competition between the remaining lines led to a new round of extinction. It is notable that alleles from the local variety, Atlas, show evidence for increased fitness in the population though it was only brought to California around a century and a half before the start of the experiment <u>(*18*)</u>.

## The emergence of a dominant clonal lineage in CCII

The paucity of genetic diversity found in generation F58 suggested that relatively few genetic lineages may have survived into the later stages of CCII. To understand the composition of the genomes of individual progeny, we conducted a genotyping by sequencing (GBS) experiment on CCII parents and 878 total progeny sampled across generations F18, F28, F50, and F58. After alignment, SNP calling, and filtering, we identified 263,238 sites suitable for further analysis.

Consistent with large-scale loss of genetic diversity in CCII over time, we find that the average genome-wide genetic distance between lines fell dramatically in later generations (Fig. 3a, Fig. S9). After hierarchical clustering of lines based on genetic distance, we found nearly identical whole genome haplotypes at all time points that have risen to high enough frequency (from an initial 1/30000) to be sampled multiple times (Fig. 3b).

**Fig. 3.**
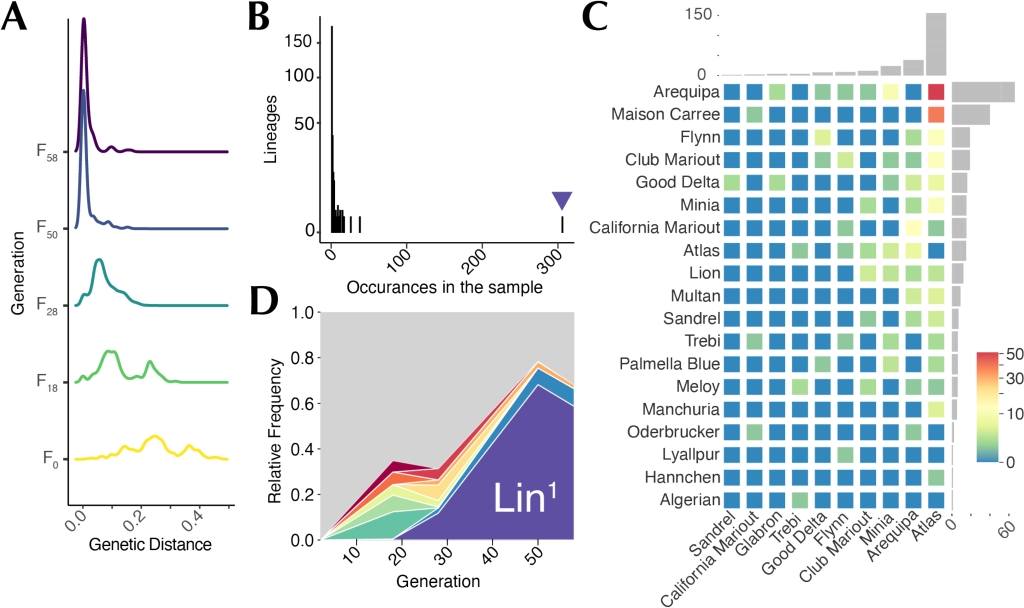
The rise of select lineages in the CCII. (**A**) Density plots of the distribution of genetic diversity across five time points in the CCII. (**B**) Nearly identical lineage abundance across the CCII. The purple arrowhead marks Lin^1^. (**C**) Primary (x axis) and secondary (y axis) ancestry for the CCII progeny. The adjacent barplots indicate the sum of individuals that show a primary or secondary relationship with each observed parent, and the colors of the heatmap indicate the number of individuals with ancestry from a particular combination of parents. (**D**) Muller plot showing the proportion of sampled individuals belonging to the top ten most common lineages over time. The grey portion of the plot are all other lineages.

Of the 878 sequenced progeny across all four generations, we identified only 261 distinct lineages by these criteria. Of these, 177 (68%) occurred just once in our sample. The remainder occurred more than once, with most individuals (445) belonging to just 8 clusters. One lineage was found at exceptionally high frequency, making up 34.9% of all sampled individuals (Fig. 3b, Lin^1^).

Principal component analysis of representative samples from each multilocus haplotype cluster revealed a close relationship between the most successful clusters and North African-derived parents (Fig. S9). To understand which specific parents contributed to successful lineages over time, we inferred the dominant ancestry of each multilocus haplotype. We assigned each lineage a primary and secondary parent defined as those which shared the greatest and second greatest fraction of the line’s genome in near identity (*d* < 0.01). Only 10 of the 28 parents were assigned a primary relationship with at least one lineage, and 19 were assigned a secondary relationship (Fig. 3c). Six of the eight primary parents identified in this analysis came from the North African group. All nine North African varieties were identified as secondary parents of at least one lineage. The Californian historical variety, Atlas, was the primary relationship in 59.7% of lineages, a massive enrichment from the expected 3.6% (Chi-square test, p < 1e-46). Potential Atlas / Arequipa progeny were the most common (19.5%), with Atlas / Maison Carre being a close second (14.9%). The most abundant lineage, Lin^1^, fell into this latter group (Fig. S10). These patterns confirm that a relatively limited number of lineages, predominantly derived from Mediterranean-adapted parents, were able to succeed in CCII.

The abundances of each of the 261 lineages were not static over evolutionary time. The number of sampled lineages decreased in later generations (93 in F18, 93 in F28, 38 in F50, and 55 in F58). This drop in diversity was largely the result of a dramatic increase in the abundance of Lin^1^ over time, rising from a frequency of 0.005 in F18 to a frequency over 0.586 in our sample of generations F50 and F58 (Fig. 3c). The runaway success of Lin^1^ appears to be driving out most of the diversity in CCII putting the population on a path to genetic homogeneity within the coming decades.

## The targets of natural selection in CCII

We next sought to pinpoint the genes targeted by directional selection during rapid adaptation in CCII. Strong selection at a locus can leave a footprint of depleted genetic diversity and strong allele frequency change surrounding the targeted site relative to the rest of the genome <u>(*19*)</u>. Because genetic variation in CCII had become so depleted by F58, we focused on identifying selected loci in the first phase of the experiment by comparing F2 and F18.

For 25,844 sliding windows (1,000 SNP windows, 500 SNP step size) covering the barley genome, we calculated the combined probability of the observed He and mean change in allele frequency (between F2 and F18) in a set of 100,000 simulations of neutral evolution (see Materials and Methods). After false discovery rate correction, we identified 58 genomic regions that showed significant evidence for selection in the population (Fig. 4, Fig S11-14). These regions ranged from 162 kb to over 42 Mb in size, with a median size just under 1 Mb (928.691 kb, Table S2), covering a total of 3.5% of the barley genome.

**Fig. 4.**
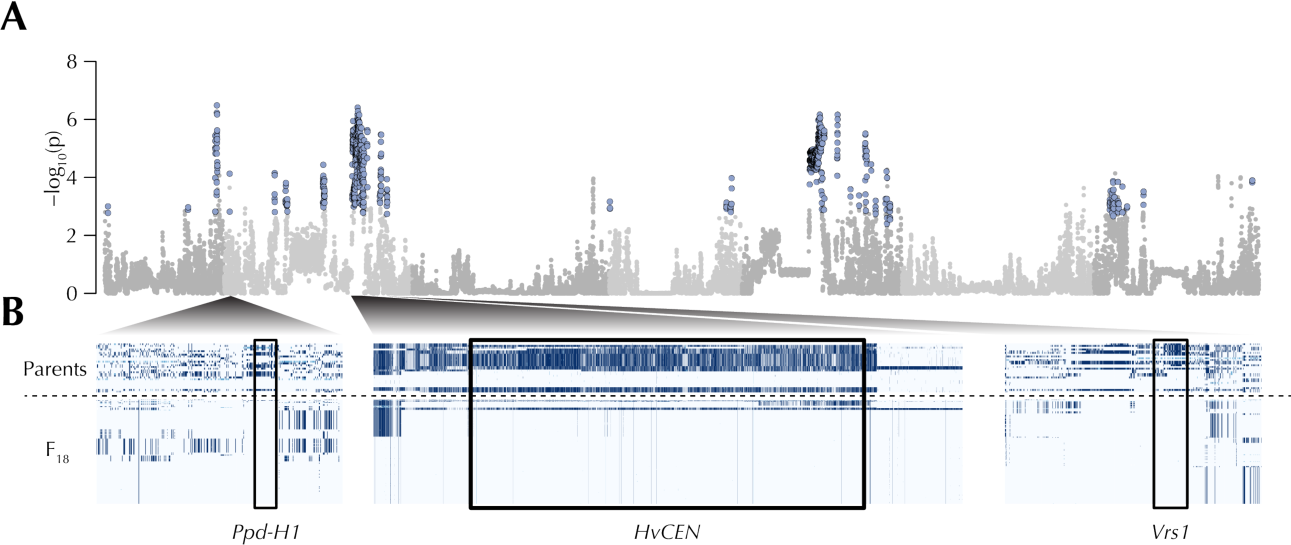
The footprint of selection in the CCII. (**A**)Manhattan plot showing significant selected regions (light blue points). (**B**)Highlighted selected regions with significant region shown as a box. Parental haplotypes are shown in the top row and F18 in the lower row. For visualization purposes SNPs were polarized against the Atlas genotype across the region so that lighter colors indicate similarity to a north African parent.

Several compelling candidates overlapped with regions of the genome targeted by natural selection. *Vrs1*, a homeobox gene which controls a major dimorphism in inflorescence architecture in barley <u>(*20*)</u>, overlapped with a 2.06 Mb selected region on the long arm of Chromosome 2H (Fig. 4, Table S2, and Fig. S6). Wild and some cultivated barley have inflorescences that produce two rows of seeds (two-rowed), but plants that carry loss of function mutations of the *Vrs1* gene produce up to six rows of seeds (six-rowed or intermedium types). 7 of the 28 CCII parents carried two-rowed alleles (Table 1, 4 from Europe, 1 from North Africa, and 2 hybrids), but by F18 the population is fixed for the six-rowed allele *vrs1.a1* in our pooled sequencing data. This allele contains a single base pair deletion that generates a frameshift in the last exon of *Vrs1*. Ultimately, the success of this allele led to extinction of the two-rowed phenotype in the population <u>(*13, 14*)</u>.

The frequency of six-rowed types in cultivated barley is often attributed to artificial selection by early farmers <u>(*20*)</u>. However, these results indicate that natural selection at Davis for the six-rowed type is likely very strong, perhaps because of the larger number of seed produced per head. Several other genes overlapping selected regions had homology to factors that affect floral development in plants (Table S2). These included genes characterized in rice (*OsLa, OsMADS22, OsLBD12, OsLG3, MINI ZINC FINGER2, Shattering1*, and *ABERRANT PANICLE ORGANIZATION1*) and in the plant model system *A. thaliana* (*AINTEGUMENTA, KRYPTONITE2*, and *UNICORN*) pointing to inflorescence development as an important target of selection over the course of the experiment <u>(*21–30*)</u>.

We also found that the signal of selection overlapped with genes that play a role in the timing of reproduction, a key adaptive trait in barley<u>(*31*)</u>. The most well-studied of these are the pseudo-response regulator gene *PHOTOPERIOD1* (*Ppd-H1*) and the phosphatidyl ethanolamine-binding protein encoding gene *HvCENTRORADIALIS* (*HvCEN*). *Ppd-H1* and *HvCEN* regulate the timing of flowering via the day-length sensing and autonomous genetic pathways respectively <u>(*32, 33*)</u>. Early flowering ancestral alleles were nearly fixed at both loci in F18 (Fig. 4b and Fig. S14 and S15). The late flowering phenotype of derived alleles of *HvCEN* is thought to result from a Proline to Alanine change at position 135 in the protein sequence. The late flowering allele at HvCEN was found at intermediate frequencies in the parents (12/28), but pooled sequencing data found only limited evidence of its continued segregation in the progeny (1/39 in F18, 1/17 in F28, and 0/21 in F58). Similarly, a late flowering allele at *Ppd-H1* was found in 8/28 parents but was no longer detected in the later generations. It should be noted that an allelic series at *Ppd-H1* regulating flowering time with differing effect sizes has been proposed, though in our dataset it seems that just one allele predominates<u>(*34*)</u>. Other selected regions overlapped with homologs of genes involved in flowering time in other species, including *OsLF* and *Hd3a BINDING REPRESSOR FACTOR2* (*HBF2*) <u>(*35, 36*)</u>.

Two unusually large, selected regions were found on chromosome 2H and chromosome 5H (Table S2 and Fig. S13). The first overlapped with *HvCEN* and a known large inversion polymorphism that segregates in barley <u>(*37*)</u>. The exceptionally large footprint we identify in our analysis (29.2 Mb) is likely the result of suppressed recombination from the inversion. The largest selected region (42.9 Mb) was found in the pericentromere of chromosome 5H. We did not identify a previously characterized candidate gene in this region, but it colocalizes with a region that appears to have been targeted by strong selection in barley breeding lines over the course of the last century <u>(*38*)</u>. It remains to be determined whether the strong signal of local adaptation in these regions might be driven by more than one gene.

Notably, the selected regions identified here have become nearly fixed in just a few decades without conscious human-mediated selection. This suggests that adaptive alleles that emerged during early agriculture could dominate a locale during a farmer’s lifetime.

## The genetic basis of stabilizing selection in CCII

We next asked how genetic shifts in CCII translated into adaptive phenotypic change. We focused on the timing of reproduction because of its general importance in plant fitness<u>(*39*)</u> and because our selection scans uncovered several candidate genes associated with this process.

The appropriate timing of reproduction is a critical contributor to barley local adaptation <u>(*40*)</u>. Flowering too early can expose delicate inflorescences to winter frosts and limits the accumulation of vegetative biomass. However, late flowering risks failure to set seeds before the onset of the hot, dry summer season. Previous work with CCII indicated that these combined forces favor intermediate flowering times, a process known as stabilizing selection <u>(*12, 15, 41*)</u>.

We confirmed these results by observing CCII progeny from F18, F28, F50, and F58 alongside the 28 parents in our greenhouses. Median flowering time (as estimated by days to awn emergence) fell modestly from the midparent values by 5.6 days (Mann-Whitney test, p < 1e-7) over the first 18 generations with a notable reduction in variance due to the extinction of late flowering outliers (F test, p < 0.001, Fig. 5a). After generation F18, flowering time slowly rose to just above the founding mean (+ 1.4 days comparing F2 to F58, + 7 days F18 to F58, Mann-Whitney test *p* < 1e-6) with greatly reduced variance (F test, *p* < 1e-13, Fig. 5). In our experiments with continued watering throughout the plant’s lifetime, flowering time was positively correlated with fitness-related traits including plant height and fecundity (Fig. 5b and Supplementary figures 17 and 18, *p* = 3.5e-56, adjusted R^2^ = 0.62). Despite the advantages of late flowering, the population did not favor the latest flowering genotypes, but instead the dominant Lin^1^ showed an intermediate flowering time (Supplementary figure 19). Together these observations indicate that flowering time in CCII is the target of stabilizing selection, which drives populations toward intermediate phenotypes by removing phenotypic extremes. An initial shift in phenotypic mean is predicted by theory if the trait distribution is skewed <u>(*42*)</u> as is the case here, but stabilizing selection on the timing of reproduction only becomes apparent in CCII on longer time scales <u>(*41*)</u>.

**Fig. 5.**
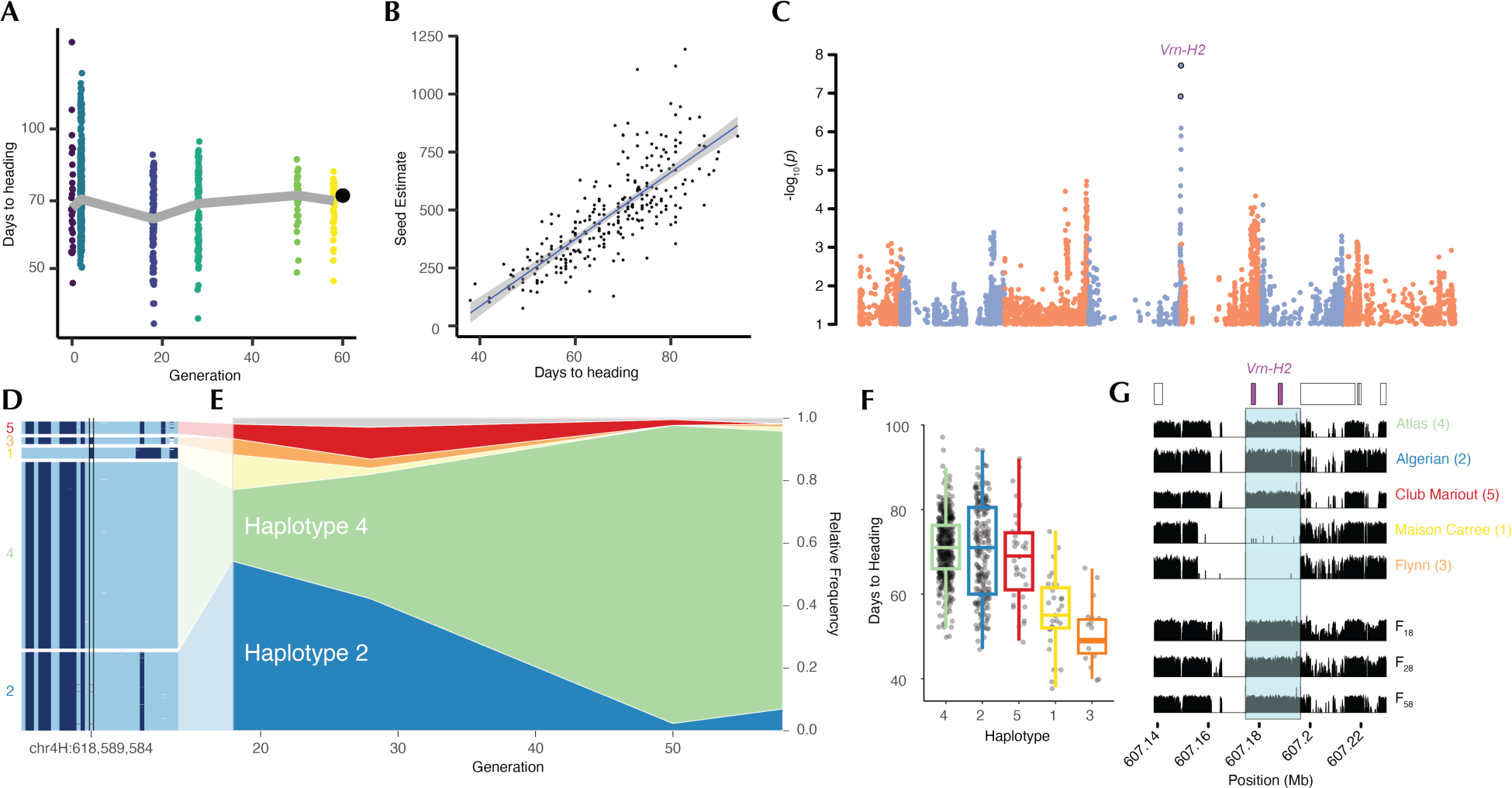
The dynamic process of purifying selection on reproductive timing. (**A**) Evolution of heading date in the CCII. The F2 distribution was estimated from midparent means (teal). The black dot shows the value Lin^1^. (**B**) The relationship between days to heading and fecundity based on seed number estimates. (**C**) Genome wide association study of days to heading showing a single significant peak (Bonferroni corrected p < 0.05) on Chr4H. (**D**) Haplotype structure surrounding the Vrn-H2 locus. The highlighted SNP is the nearest significant SNP to the *Vrn-H2* deletion. (**E**) Muller plot showing haplotype frequency from generation 18 to 58. (**F**) Distribution of days to heading across the five main haplotypes segregating in the CCII. (**G**) Coverage of sequencing reads across the *Vrn-H2* locus in the B1K-04-12 *H. spontaneum* assembly for examples of each haplotype group. The highlighted region (light blue) shows the segregating deletion which overlaps with two annotated ZCCT genes. The three bottom tracks show coverage for each progeny pool.

The success of alleles derived from Mediterranean-adapted parents at *Ppd-H1* and *HvCEN* is consistent with selection against late flowering types in CCII (Fig. 4 and Supplementary figures 14 and 15). This left us to explore how stabilizing selection could drive a second shift by eliminating the earliest flowering progeny. We conducted a genome wide association study (GWAS) to discover loci associated with variation in flowering time and identified a single significant peak on the long arm of chromosome 4H (Fig. 5C), which we verified using a subset of lines in the following year (Supplemental figure 20). The lead SNP in our analysis was in linkage disequilibrium with a well-characterized regulator of adaptive shifts in barley flowering time, *VERNALIZATION2* (*Vrn-H2*, the tag SNP is 5.42 Mb from the locus and in LD, r^2^ = 0.83, with the closest significant SNP to the locus 53kb from the reported causal lesion).

The ancestral barley *Vrn-H2* locus encodes three zinc-finger/CCT domain (ZCCT) transcription factors that repress flowering in the absence of prolonged cold treatment, preventing early flowering during the winter <u>(*43, 44*)</u>. A large deletion common in northern European barley spans all three genes and drives early flowering in the absence of extended cold exposure. CCII segregated for eight haplotypes at the *Vrn-H2* locus, five of which were found at an allele frequency greater than 0.05 in at least one of the sampled generations (Fig. 5D). Three North African alleles (Haplotypes 2, 4, and 5) conferred late flowering and together were found in 92.4% of the sampled progeny and 81.8% of the F18 (Fig. 5E and F). The two early flowering alleles, Haplotype 1 and 3, were less common and less geographically restricted being shared by Mediterranean and Northern European parents (16.4% of F18 and 5.9% of all sampled progeny). The late flowering Haplotype 4 from Atlas increased rapidly over time and by generation F58 88.9% of individuals carried this allele while early flowering alleles dropped to 2.3% of the sample. The rise of Haplotype 4 corresponds to an almost complete loss of the minor allele at the tag SNP in our GWAS.

By realigning the parental whole genome sequences to a reference that contained a functional ZCCT gene cluster (*H. spontaneum*, B1K-04-12; Supplementary figure 22)<u>(*37*)</u>, we found clear evidence for the presence of the ZCCT genes in the three late flowering Haplotypes 2, 4, and 5 and in the pooled sequencing samples at all time points (Fig. 5G). The two early flowering alleles lacked coverage across both genes, consistent with a deletion.

Together our results indicate that stabilizing selection tuned flowering time in CCII through a two-step process. First, selection for Mediterranean alleles at *Ppd-H1, HvCEN,* and perhaps other loci assured completion of the life cycle before the onset of the dry season. Second, selection for a functional *Vrn-H2* allele from Atlas eliminated premature flowering in the winter or early spring. The unusual length of CCII experiment made it possible to resolve the long-term trend of stabilizing selection that emerges from shorter directional shifts in flowering time.

Our analysis of the CCII reveals the power of selection to rapidly drive strong directional evolution even in a variable environment, which is consistent with recent results from field experiments with Drosophila <u>(*45*)</u>. Low genetic diversity in wild populations of self-fertilizing plants is often attributed to founder effects, though linked selection has also been proposed as a potentially important driver <u>(*46*)</u>. The latter appears to be the most important force in the CCII, with high genetic diversity in the founding population rapidly obscured by the pervasive effects of natural selection. The footprint of early selective events has already begun to decay after only a few dozen generations, suggesting that historical selection may be obscured when divergences are larger than a few tens of generations. However, the length of the CCII experiment allowed us to observe the dynamics of asymmetric stabilizing selection, which over shorter periods would have been disguised as directional selection. These findings highlight the challenge of interpreting patterns of genetic diversity in large multicellular organisms; long-term observation is difficult but crucial. An improved model of the dynamics of adaptive evolution based on observation has great potential to facilitate prediction of patterns of genetic diversity and to aid in the development of strategies for countering the impacts of rapidly changing environments.

## Acknowledgments

We would like to thank the leadership and staff at the UCR Agricultural Experimental Station, the leadership staff of the UCR High-Performance Computing Center, and Harold Bockelman at the USDA Small Grains and Potato Germplasm Research program for supporting the infrastructure and resources necessary to complete this project. We thank Cal Qualset for providing the composite cross germplasm and for comments on the manuscript. We would also like to thank members of the Koenig and Seymour labs for their comments and suggestions during the conduct of this research. We acknowledge the assistance of Felipe Reyes Gaibor in the initial curation of prior publications on CCII populations and the current availability of parental seed stock. Finally, we would like to thank the researchers including Harry Halan, Mary Martini, Coit Suneson, and Gus Wiebe, who worked to maintain composite cross experiments over the last one hundred years.

## Author contributions

Conceptualization: DK Methodology: DK, JBL, PLM

Formal Analysis: DK, JBL, KEB, CJF Investigation: JBL, AMG

Funding acquisition: DK, JBL, KEB Project administration: DK, JBL Supervision: DK

Writing – original draft: DK

Writing – review & editing: DK, JBL, KEB, CJF, AMG, PLM

## Data and materials availability

Genotype by sequencing datasets have been deposited at the National Center for Biotechnology Information (NCBI) under BioProject accessions PRJNA575963. Whole genome sequencing datasets have been deposited at the National Center for Biotechnology Information (NCBI) under BioProject accession PRJNA1018527.

## Supplementary Materials

Materials and Methods

Supplementary Text

Figs. S1 to S23 Tables S1 to S2 References (##–##)

## Materials and Methods

### Plant Material

The composite Cross II experiment has been propagated every few years since its inception in population sizes of approximately 10,000-25,000 plants. Seed viability for early generations was maintained by propagating less frequently, for example every five years rather than every year (Fig. 1). The population was maintained with minimal human intervention excepting planting in the fall and harvest in the spring. See Supplementary Material for further details.

### <u>CCII parent whole genome</u> sequencing

Nuclear genomic DNA was extracted using the CTAB method (47) independently from two plants each of the 28 parents of CCII (48). The first replicate of DNA extractions was used as input for the Illumina TruSeq DNA PCR-Free library preparation kit. Libraries were each sequenced on a single lane of an Illumina HiSeq 4000 instrument on the paired-end 150 bp read setting. To increase depth of coverage across the parents, libraries were generated from the second replicate using the Nextera DNA Flex library preparation kit. These libraries were pooled and sequenced on an Illumina NovaSeq instrument to generated paired-end 150 bp reads.

### Pooled DNA extractions and genome sequencing

One thousand seed from generations F18, F28, and F58 were planted in flats and grown until the two-leaf stage. A hole punch of leaf tissue was taken from each plant and combined together across all plants to generate a single pool of tissue that was then finally ground in liquid nitrogen to homogeneity with a mortar and pestle. A subsample was then taken and used as input for CTAB genomic DNA extraction and sheared to 300 bp fragments on a Covaris S220 Ultrasonicator. A library for each generation was constructed with the NEB Nextseq kit library preparation kit. The three libraries were independently pooled four times and run on four lanes of an Illumina HiSeq 4000. To increase coverage, the parents were resampled and extracted DNA was used to generate sequencing libraries with the Illumina Nextera Flex library preparation kit pooled and sequenced on an Illumina Novaseq instrument.

### Sequence alignment and variant calling for CCII parents

The parental and progeny pool DNA sequences were aligned to the Morex 2019 TRITEX barley reference genome assembly (also referred to as v.2)(49) using the bwa mem short read alignment algorithm v. 0.7.17-r1188 (50) and sorted using samtools v. 1.14 (51). Sorted bam files were input into the standard pipeline to identify single nucleotide polymorphisms using GATK v. 4.1.4.1 (52). Initial alignments were screened for PCR duplications using the markduplicates tool in GATK, with the subsequent alignment files being used one chromosome and sample at a time as input for the GATK haplotypecaller tool. The CombineGVCFs tool was used to combine data across parents, and then variant calls were made using the GenotypeGVCFs tool allowing a maximum of two alternate alleles at variant sites. The raw variant dataset from each chromosome was filtered using vcftools v. 0.1.17 (53) and bcftools v. 1.15 (54) to include only SNPs with no more than 4 parents having missing data, a minimum genotype quality of 5, a variant quality of 30, a maximum of 2 alleles, no more than 6 heterozygous calls, and a minimum per sample depth of 2. We also required at least one observation of an individual homozygous for the alternate allele. The bcftools concat tool was then used to combine the multiple chromosomes of barley into a single file. Variant sites were annotated using SNPeff (v.5.1)(55).

### Exome capture variant calling and analysis with CCII parents

Raw Illumina read data from the barley exome capture panel (10) were downloaded from the European Nucleotide Archive and aligned with bwa mem to the Morex 2019 Tritex assembly (49). Adapter sequences were trimmed using trim_galore wrapper for cutadapt (v0.6.7)(56). Variant calling proceeded according to the standard practices recommended for the GATK tool as described for the whole genome sequencing for CCII parents. SNP filtering was performed using vcftools v.0.1.16-18 (53) with the options: --max-missing 0.8 --maf 0.01 --mac 3 --minDP 10 --maxDP 500 --minQ 10 --min-alleles 2 --max-alleles 2.

Genotype calls for each parental accession were made at each polymorphic site in the exome panel using the samtools mpileup and call tools (54) and merged with the exome dataset using bcftools merge. The final dataset was filtered to include only domesticated samples from the exome capture dataset, leaving 1,739,465 biallelic SNPs.

Principal component analysis of the combined sample was conducted with the software PLINK v1.90b6.25 (57) considering the first 25 axes of variation. Plotting and analysis of allele frequencies was performed using the R statistical computing environment (58).

### Pooled sequencing allele frequency and diversity estimations

Alignments for pooled sequencing samples were conducted as described for parental genome sequences. Allele counts at each of the sites polymorphic in the parents were generated for each alignment file using the script extractsite_counts.py which directly extracts the observed nucleotides at each position from the alignment. Count data were then summed across bam files to generate a final allele count dataset for F18, F28, and F58, which was then combined with allele counts from the parents generated from the filtered VCF file using the script Process_Filtered.py. A final filter was applied to keep sites with a coverage of < 300 across all samples and > 10 counts for each generation in the R statistical computing environment (58).

Allele frequencies for alleles p and q in population n at site l were calculated from the count data as follows:

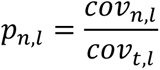

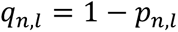

Where #$%_$_ is the total coverage of the site, and #$%_!_ is the coverage of the first allele at the site. For genome wide expected heterozygosity calculation and simulation, only sites with complete data across the parents were considered. Expected heterozygosity for a specific population was calculated at each site was calculated as follows:

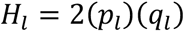

Where *H_1_* is the estimate of expected heterozygosity at site l. For windowed of genome wide estimates:

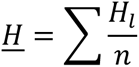

Where n is the total number of segregating sites in the window. Simulations of loss of genome-wide ) over time were conducted under the conservative assumption of no recombination and complete selfing. An initial population of the specified size seeded the simulation with each individual representing a parental genome in equal proportion. Subsequent generations were derived from a random sample of the previous generation with replacement. Expected heterozygosity was calculated for generations F18, F28, and F58 as above, using the summed allele counts of the remaining genomes.

Genome-wide pairwise Wright’s Fst was calculated as:

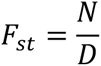

where

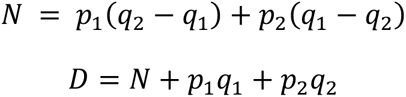

With n being the specific pool. The mean across sites was used to describe genome-wide *F_ST_*

To understand the relationships between the population and the parents over time, we calculated Nei’s genetic distance (59). Let *X_u_* be the frequency of allele u in population X at site l. The distance D between each population was calculated as follows:

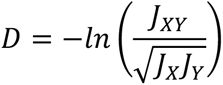

where;

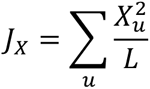

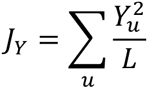

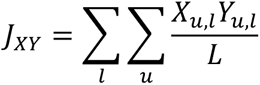

The resulting distance matrix was used to conduct a multidimensional scaling analysis implemented in R.

### Realignment of the whole genome sequences against the *H. spontaneum reference*

The Morex reference contains the null, deletion allele at *Vrn-H2* preventing us from determining segregation of functional alleles in the parents. For this purpose, the parental and pooled sequencing reads were realigned against the *H. spontaneum* reference B1K-04-12(5), which contains two of the three characterized ZCCT genes. The sequencing depth was calculated per base pair using samtools v. 1.14 for primary read mappings of a minimum mapping quality of 60. The reported coverage data shows 21 bp sliding medians of these values calculated and plotted in R.

### Selection scans

We scanned the genome in 1,000 SNP windows for signals of selection between generations F0 and F18using the combined signals of change in allele frequency (Δp) and loss of H. We began by simulating neutral evolution of each window using the known starting allele frequencies. The simulated population sizes were held at the very conservative size of N = 100 to account for the genome-wide loss of genetic diversity due to LD. Coverage variability was simulated for each site as normally distributed, with a mean of 25 and variance of 7.2. Evolution was then simulated for 16 generations (equivalent to F18) for each window, like the whole genome simulations described above. At the final time point, the mean Δp and He were calculated and recorded. 100,000 simulations were conducted to generate the distribution of both statistics expected under a neutral model.

A one-sided p-value was then calculated for each statistic in each window based on the likelihood of the observed value in the simulated data, and Fisher’s method was used to generate a combined p-value for both tests. Significant regions were defined as those which showed p < 0.05 after multiple testing correction using the Benjamini-Hochberg Procedure. Regions were additionally required to have p < 0.05 for both He and Δp tests without multiple testing correction. Adjacent windows were collapsed into single regions for reporting.

For Vrs1, Ppd-h1, HvCEN we directly scored the frequency of previously reported causal mutations. These were not necessarily included in our parental variant panel because of stringent filtering or because we excluded deletions from this set. We directly observed the frequency alleles across generations in our alignment files using the Integrative genomics viewer (60).

### Genotype by sequencing dataset

A random selection of 220 seeds from each of F18, F28, F50, and F58 and the parents were surface sterilized and stratified for 1 week on wet paper towels before germination on the bench top. Individual seedlings were transplanted into pots in the greenhouse within a week of germination. Young leaf tissue from each progeny and three replicates of each parent was harvested and transferred into 96 well plates. CTAB genomic DNA extractions were performed for each sample, followed by DNA quantification and quality assessment on agarose gels and using the Quant-It PicoGreen assay (Thermofisher, P11496). Sequencing library construction followed (61). Sequencing was performed on an Illumina HiSeq 4000 instrument using single-end PE 150bp run mode.

Reads for the parents were parsed by barcode, trimmed for adaptor sequences and low-quality ends using the software package trimmomatic v.0.39(62), and aligned to the TRITEX barley reference genome (49) using the minimap2 software v.2.1 (63). Alignments were sorted and aligned using samtools v.1.17 (51). Polymorphic sites were identified using bcftools v.1.17 (54) using the mpileup and call tools with the flags -c -v. bcftools was then used to filter the raw calls to include only SNPs with < 4 heterozygous calls, a minimum coverage of 3 reads in each parent, a total depth < 100 but less than 1000 across all parents, a quality of more than 500 and no more than 1 alternate allele.

Trimming and alignment was conducted for the progeny as described for the parents. SNP calling at each parental polymorphism in the progeny was conducted using the approach implemented in STITCH (64) using a K = 28 and nGen = 60. Imputed genotype data were then merged across chromosomes and filtered with bcftools for sites with missing data < 0.05 and < 5 heterozygous calls.

Genetic distance calculation and principal component analysis between progeny and parent datasets were performed using PLINK v.1.90b6.25 (57). Genetic distances of progeny lines were used as input for hierarchical clustering with average linkage and a tree cut with a 0.001 threshold. This approach was used to select a random member of each cluster for genome wide association studies. A consensus haplotype was also generated for ancestry estimation based on the majority rule at each site. The consensus was then used as input for ancestry calculations.

Assignment of primary and secondary ancestry was accomplished using a windowed analysis. For each progeny, the total number of 1 Mb genomic windows falling below a genetic distance threshold of 0.01 from a parent were recorded. The parent with the largest fraction of windows falling below this threshold was assigned as the primary parent. The secondary parent was assigned as the parent with the largest fraction of remaining windows falling under the 0.01 threshold. This approach is approximate because of allele sharing between the parents, segregating variation within the parent accession not captured in our resequencing samples, and several near identical parents.

### <u>Greenhouse flowering time</u> experiment

In the fall of 2016 and 2017, seeds from the parents and progeny were surface sterilized using 15% bleach, washed with distilled water, and placed on a damp paper towel in a plastic container. Each container was stratified at 4C for one week in the dark, and then seeds were placed at room temperature on the benchtop for four days to allow germination. Individual seedlings were transplanted into 1L pots containing UC Soil mix 3 in a greenhouse in Riverside, CA, USA set to maintain temperatures above 20C and with artificial light set for 14:10 long day conditions in a completely randomized design. In 2016, four replicates of each parent were transplanted, with one individual each of the progeny genotypes. In 2017, a subset of progeny genotypes were selected based on pruning of nearly identical samples and two replicates per line were planted in a completely randomized design. Flowering time was estimated based on the emergence of awns (awn tipping) from the first inflorescence. Plant height was measured after plant dry down during harvest. An estimate of the total seed number, Ŝ, was generated with the following formula:

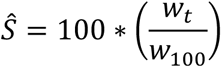

Where w_t_ is total seed weight and *w*_100_ is hundred seed weight.

When available, replicate data were summarized for both parents and progeny as the mean of observed values.

### Genome-wide Association Studies

The down sampled progeny dataset, which contained a single member drawn at random from each of the near clonal genetic lineages, was used to conduct a genome-wide association study on heading date. GWAS were conducted using the software gemma v.0.98.1 (65) with cryptic relatedness correction using a kinship matrix calculated from the subsampled genetic dataset.

## Supplementary Text

### CCII parental diversity and founding relationships

The parents of the Composite Cross II were selected to represent the global diversity of cultivated barley at the time. Harlan assigned twenty three parents to ten geographic regions with slight differences between publications (*1,14*): five lines from northwest Africa (Algerian, Arequipa, Atlas, Maison Carré, and Sandrel); four from Egypt (California Mariout, Club Mariout, Good Delta, and Minia); four from north Eurasia (Hannchen, Horn, Manchuria, and Oderbrucker); two each from India (Lyllapur and Multan), the Soviet Union (Orel and Lion), the eastern Mediterranean (Palmella Blue and White Smyrna); and finally one each from Armenia (Trebi), China (Han River), the Balkans (Wisconsin Winter), and Mount Everest (Everest). Of these, Arequipa, California Mariout, Everest, Han River, Good Delta, Minia, Orel, and Palmela Blue are unselected traditional varieties with the remaining accessions being pureline selections from traditional varieties. The two Indian accessions are of unknown improvement status (Table S1). The parents were phenotypically diverse with a variety of grain colors; 21 six-row and 7 two row varieties; and 22 rough awned, 3 smooth, and 2 semi-smooth varieties.

We filtered variants amongst CCII parents to include those segregating above a minor allele frequency threshold of 0.05 and having low levels of local linkage disequilibrium using PLINK v1.90b6.25 3 --indep-pairwise 100 20 .5. This left 588,248 segregating sites which we then used to conduct a principal component analysis of genetic variation amongst the parents (Fig. S3). The first principal component separated the south Asian accessions Lyllapur and Multan from all other accessions. The Nepalese accession Everest was found in an intermediate position on PC 1. PC2 separated northern European from six-rowed Mediterranean accessions. The observed population structure of the parents is consistent with population structure previously observed in larger genotyped panels (*6,10*).

Most of the relationships between parents described by Harlan were mirrored in our analysis as well. All the Mediterranean parents clustered in PCA space except for Sandrel, although the Egyptian and North African groups were not well separated from each other. Harlan previously noted that Sandrel’s origin was unclear, having been acquired from a seed seller in Europe. Atlas and Arequipa’s were collected in North and South America respectively, but their assignment to the North African group was based on historical information that lines of this type were grown in Spain. Palmella Blue is listed by Harlan as being collected in Palestine, however inspection of the import documentation that is associated with this line lists it as a pureline selection from a field collection in the Mariout region of Egypt. For this reason, the Palmella is treated as Egyptian in this study.

The northern European accessions separated from the Mediterranean barley in PC space, though the internal structure amongst this group was not easily interpreted. The two rowed and six rowed barleys were not distinct on the first two PCs, but we still separate two and six rowed European accessions for the purpose of this study because two row phenotypes are clearly maladaptive in CCII (*13,14*). Five hybrid lines were also used as parents (Flynn, Glabron, Golden Pheasant, Meloy, and Alpha), some of which were suspected or known to share ancestry with other parents (Alpha with Manchuria, Flynn with Club Mariout and Lion, Glabron with Lion and Manchuria, and Meloy with Atlas). These four hybrids were found in proximity to the described parents in PC space, and the fifth, Golden Pheasant, clustered with other two row Northern European varieties.

A few accessions are notably similar to one another, sharing identity by state at more than 90% of the genotyped sites. Of these, Club Mariout and Minia are both described as representative of the ancient variety grown in the Egyptian Mariout region, with Minia being an apparent resampling. Lyllapur and Multan were both from the same Indian seedbank and represent very similar varieties. The similarity between Sandrel and Trebi is most surprising based on their origin as described in the literature, though the true origin of Sandrel is a mystery, and it is plausible that it may have originally come from Turkey.

Using this information, we divided the parents of CCII into five main groups: Asian, European two-row, European six-row, Mediterranean (which is almost entirely composed of North African samples), and hybrid.

### A note on the initial crossing scheme and initial allele frequencies

The initial crossing scheme attempted all intercrosses in one direction. However, Arequipa x Good Delta did not take, and Wisconsin Winter x Horn and Wisconsin Winter x White Smyrna were included twice. This resulted in 379 pooled families at the beginning of the experiment (rather than the expect 378 from a half-diallel of 28 lines). Harlan reported that these two Wisconsin Winter parents may have differed genetically (*14*).

### A note on the maintenance of CCII

The early generations of CCII were maintained by storing seed and the less frequent propagation (Fig. S1). It is important to note that this means that the early generations sampled in this study are not necessarily direct ancestors of the later generations. To our knowledge, there is no existing record of the exact propagation history of the experiment. Fig. S1 represents our best estimate of the relationship between generations, and we found that Fst divergence fits this model relatively well (Fig. S7). However, minor deviation from this scheme is possible. The analyses contained in this manuscript are robust to minor deviations in relatedness between the generations.

### The relationship between allele frequency in the parents and global population

The selection of a small number of parents to found CCII could distort allele frequencies and exclude important common genetic variation in the species. We explored this possibility using a previously generated exome capture dataset that included a sample of 137 domesticated barley sampled throughout the species’ range 4. We re-identified segregating genetic variants in the exome capture dataset and then genotyped CCII parents at each polymorphic site (Materials and Methods). Common genetic variation in barley was strikingly well represented in the dataset (Fig. S2). The allele frequency spectrum observed in the 28 parents was not strongly distorted relative to the exome dataset, and allele frequencies of specific variants were strongly correlated across the datasets (Fig. S3).

### The footprint of selection in CCII

We chose two statistics as the focus of our selection scan. First, we searched for regions of the genome that had been nearly cleared of founding genetic variation. This would suggest that the population was near fixed for a single allele. The second signal we used was absolute change in allele frequency. This selects for alleles that were present at low frequencies in the founders but increased to very high frequencies, or those which started at intermediate frequencies and were rapidly lost. These two statistics are indicators of selective sweeps that favor specific alleles segregating in the parents outcompeting all other variation at a locus.

The genome-wide distribution of these statistics changed over time (Fig. S11 and S12). Windowed He was unimodal in the parents but was bimodal in F18and F28. The genome seemed to be composed of two groups of windows, those that appeared to have lost a modest amount of variation and a second set that were nearly cleared of all genetic variation. By generation F58 nearly the entire genome fell into this second category. Allele frequencies did not change as rapidly genome-wide, with most windows experiencing a small negative change in allele frequencies due to the loss of low frequency variants. The fraction of the genome experiencing strong shifts in allele frequencies, including strong increases in the frequency of minor alleles, increased over time resulting in a bimodal distribution across windows in F58. Thus, we observe a process of diversity loss over time, ultimately resulting in a near homogenous population throughout most of the genome, with strong genome-wide shifts in allele frequencies. For our subsequent analysis, we focused on F18because it represents an immediate state along this trajectory where strongly selected regions might be detected against a background of less dramatic change. The significant regions from this analysis can be found in Table S2, Fig. S13, and Fig. 4.

We also examined the persistence of the signal of selection over time (Fig. S23). Median He was higher in selected regions in the parents than in the rest of the genome. This suggests that our approach was biased to some extent to select for high founding diversity. As such, one should consider the regions identified here as outstanding examples of selection, rather than a complete library of selected loci. The median He remained below the genome-wide background throughout the experiment, though the difference was reduced after F18. In contrast the absolute allele frequency change was persistently higher than the genomic background throughout the experiment. In F58 the genome-wide value began to increase slightly.

### Genetic diversity at candidate targets of selection

We examined three of the select loci from our analysis in detail because of they have been previously hypothesized to be targets of selection during barley diversification. The flowering time regulator *Ppd-h1* overlaps with a 357,893 bp selected region on chromosome 2H. *Ppd-h1* is an important regulator of plant response to photoperiod, specifically in response to long days (*32*). Most wild barley and many cultivated barleys are long day sensitive. This allows plants to germinate in the fall, over-winter, and then flower coinciding with the onset of long days in the spring. However, when cultivated barley expanded its range into northern Europe, it was often planted in the spring to take advantage of the wet summer season. Rapid flowering in response to long days in this environment became disadvantageous. Alleles that disrupt *Ppd-H1* function drive day length insensitivity and are common in northern European varieties. A substitution of Serine for Proline in the sixth exon of Ppd-H1 is most closely associated with day length insensitivity, though as mentioned in the text an allelic series has been proposed (*34*).

As has been observed at this locus in previous studies, many alleles segregated at the *Ppd-H1* locus in CCII parents (Fig. S14). The two most common alleles in the parents were an allele shared by 8/9 of the North African parents and an allele shared by 5 of the northern European parents and 3 hybrid parents. The northern European allele carries the derived mutation that confers late flowering. The early flowering North African allele was fixed in our sample of F18, but the sample of F28 did segregate at low frequencies for other alleles suggesting it was not completely fixed population wide until later in the experiment (Fig. S14). However, we only detected the North African allele in generations F50 and F58.

Like *Ppd-H1*, late flowering alleles of *HvCEN* are thought to have facilitated colonization of northern Europe, but through the autonomous flowering time pathway instead of by changing day-length sensitivity (*33*). Late flowering was driven by a A135P substitution in *HvCEN*. Three alleles were common in the parents. The three alleles were not as clearly geographically defined, with the early flowering allele found in 12 parents including Atlas. The early flowering allele present in rose to near fixation by F18 and stayed at this frequency throughout the experiment (Fig. S15).

The parents of CCII segregated for 2 two-row alleles (*vrs.b2*; 1 accession and *vrs1.b3*; 6 accessions) and four six-row alleles (*vrs.a1*;16 accessions, *vrs.a2*; 1 accession, and *vrs.a3*; 4 accessions) (*20*). The population was rapidly dominated by the six row allele *vrs.a1* that is found in the Morex reference genome and contains a single nucleotide deletion that truncates the VRS1 protein in the final exon (Fig. S16).

### GWAS of flowering time in 2017

A selection of genotyped progeny focused on F18were regrown in 2017 to validate the effect of *Vrn-H2* on flowering time in the population (Fig. S20). This smaller number of genotypes was grown in a completely randomized design with two replicates. In addition, to replication of the peak on chromosome 4HL, we found as second significant peak on 5HL. This peak was not near the known flowering time regulator Vrn-H1 on chromosome 5, and no known regulator of flowering time was found in this region.

### <u>Explanation of the GWAS</u> strategy

To avoid biases in GWAS that result from the large number of identical individuals in CCII and the stratification of genotypes across evolutionary time, we randomly down sampled our phenotypic measurements to include just one representative individual from each lineage and conducted GWAS on this reduced dataset. Despite the presumed reduced power of this dataset, we identified a strong association across both years of our experiment (Fig. 5 and Fig. S20).

### Segregation of the *Vrn-H2* deletion amongst CCII parents

We aligned our whole genome sequencing reads against the *H. spontaneum* reference genome11 to assess the presence of ZCCT genes at the *Vrn-H2* locus in the parents (Fig. 5 and Fig. S22).12 This reference has annotations for only 2 of the 3 ZCCT genes described in the literature, but there is a unalignable gap just upstream of the two genes that may explain the missing sequence. To our surprise, 15/28 parents showed coverage across both ZCCT genes. Three of these were the three parents listed as winter varieties (Wisconsin Winter, Han River, and Orel), but the remainder are all Spring varieties including 8 of 9 North African varieties. The lines carrying deletions of the locus were all European or hybrid. Interestingly a subtle difference in pattern of coverage was detected between lines carrying deletions, with a slightly large deletion apparent in the accessions Manchuria, Horn, Oderbrucker, and Glabron.

Colors represent the population groupings of the parents (Mediterranean, orange; north European, blue; Asian, green; and hybrid, purple). The dashed line represents Atlas.

---

**Fig. S1.**
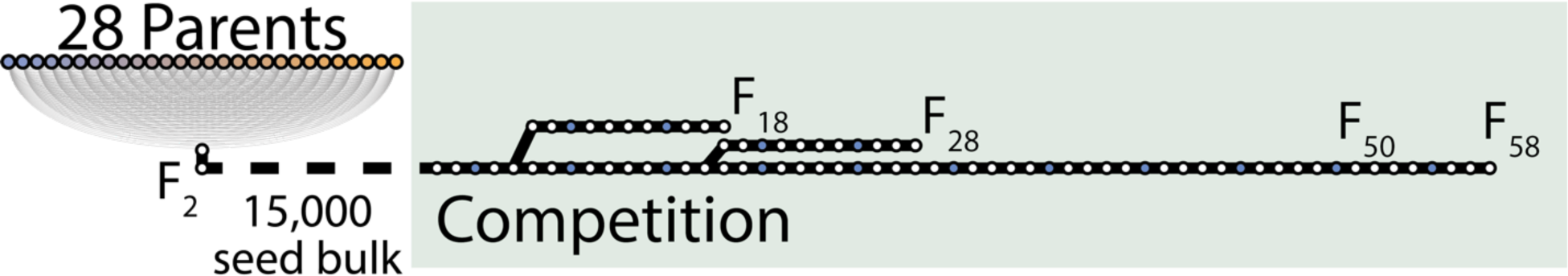
Asynchronous propagation of the composite cross II and the sampling in this study. Early generations of the composite cross were maintained by propagating some lineages less frequently than others. The relationships of the samples used in this study inferred from historical records are shown here.

**Fig. S2.**
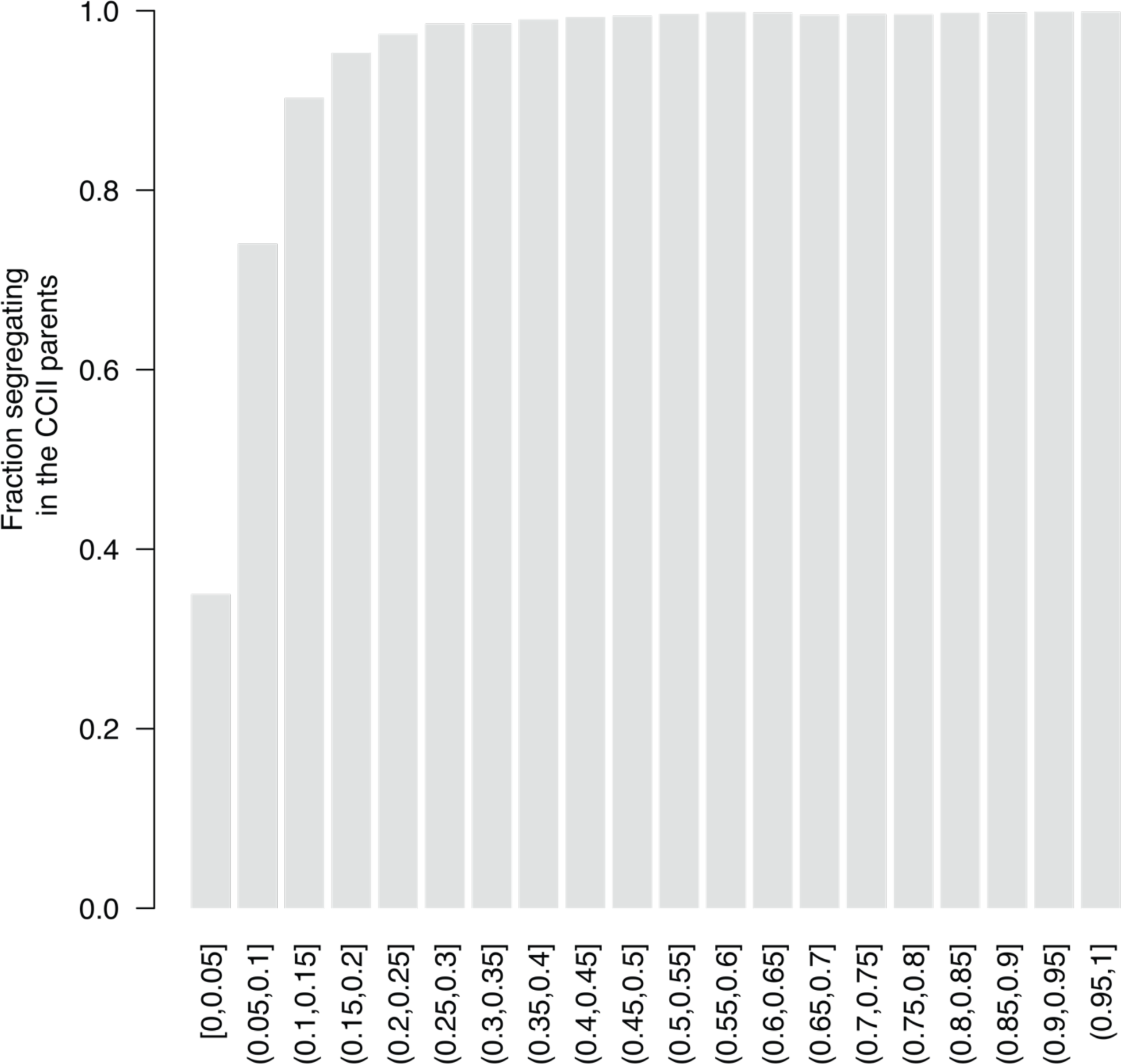
Fraction of global diversity segregating in the CCII by allele frequency.

**Fig. S3.**
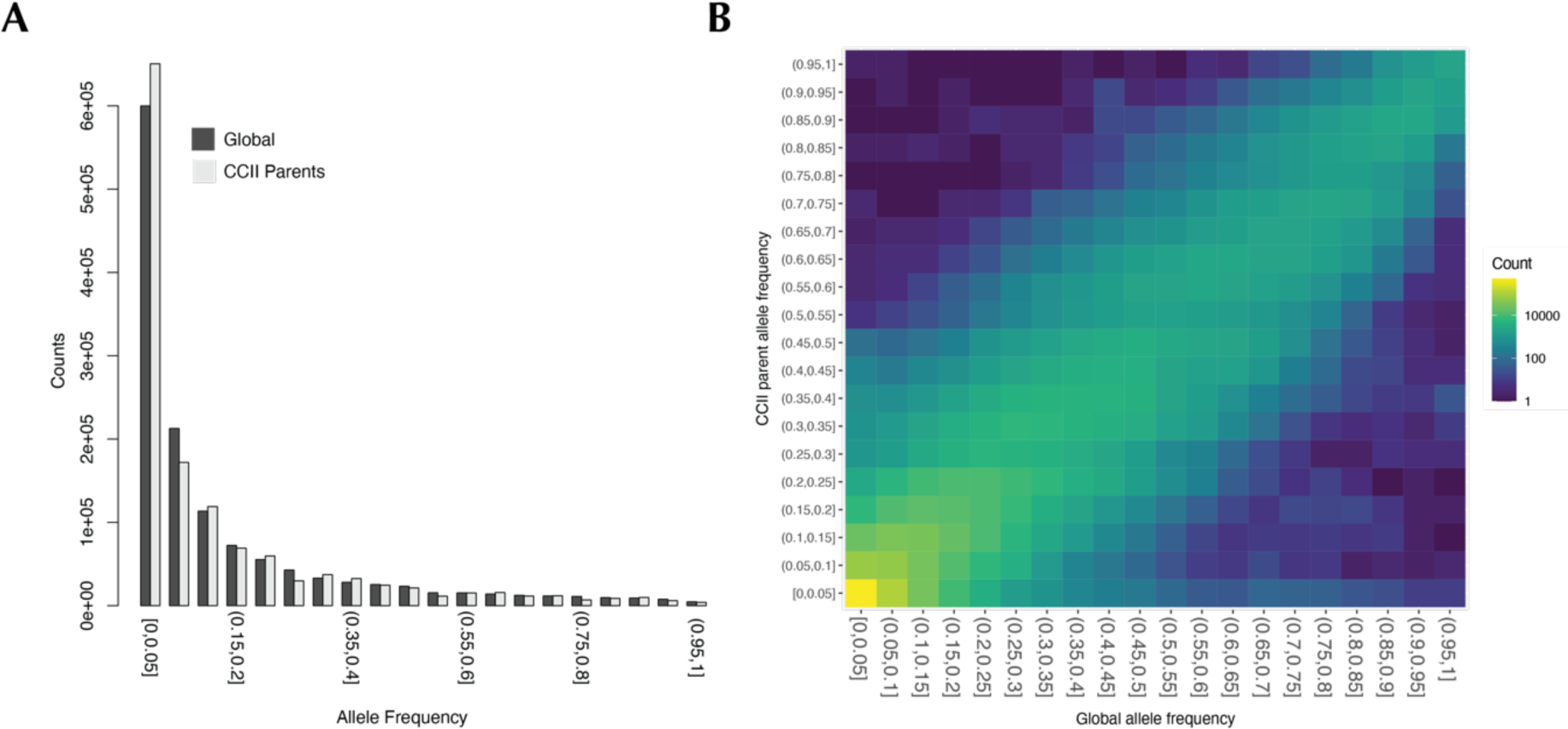
Comparison of allele frequency spectra in a global sample and the CCII parents. (**A**) the 1D allele frequency spectrum at sites segregating in the global panel. (**B**) The 2D allele frequency spectrum comparing the two datasets at sites segregating in the global panel.

**Fig. S4.**
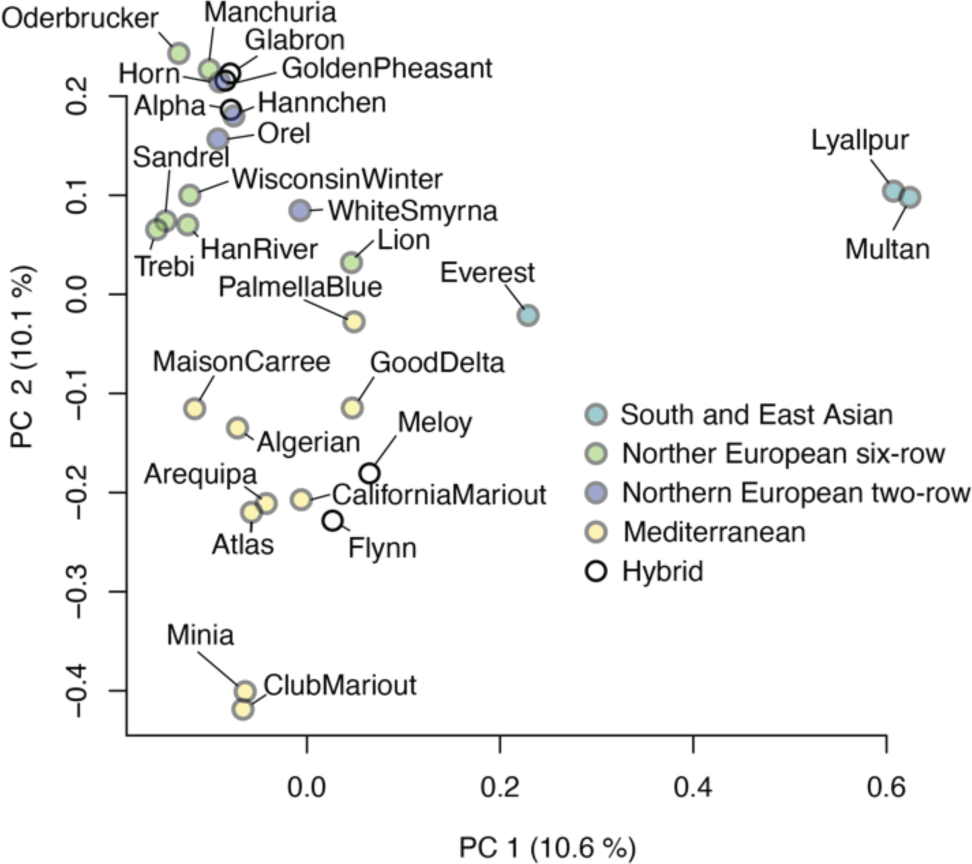
Principal component analysis of the genetic relationships between the CCII parents.

**Fig. S5.**
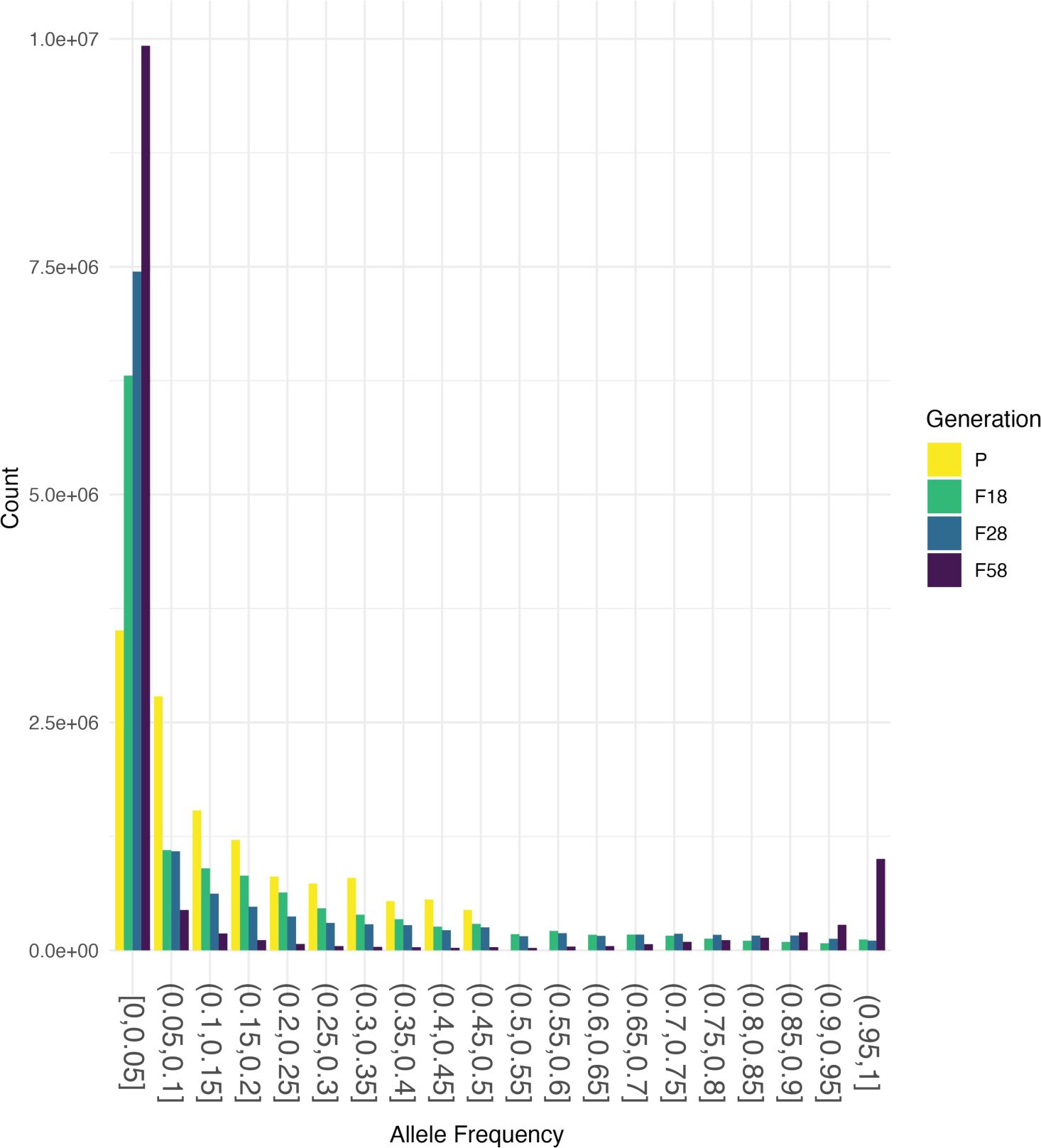
>The shift in global allele frequency spectrum over time. The allele frequency spectrum calculated for the minor allele in the parental sample across generations. The population rapidly becomes more homogeneous over time, with a loss of intermediate frequency alleles. Most of this shift is through loss of the less common allele at the founding of the experiment, although a clear subset of minor alleles has been successful over time.

**Fig. S6.**
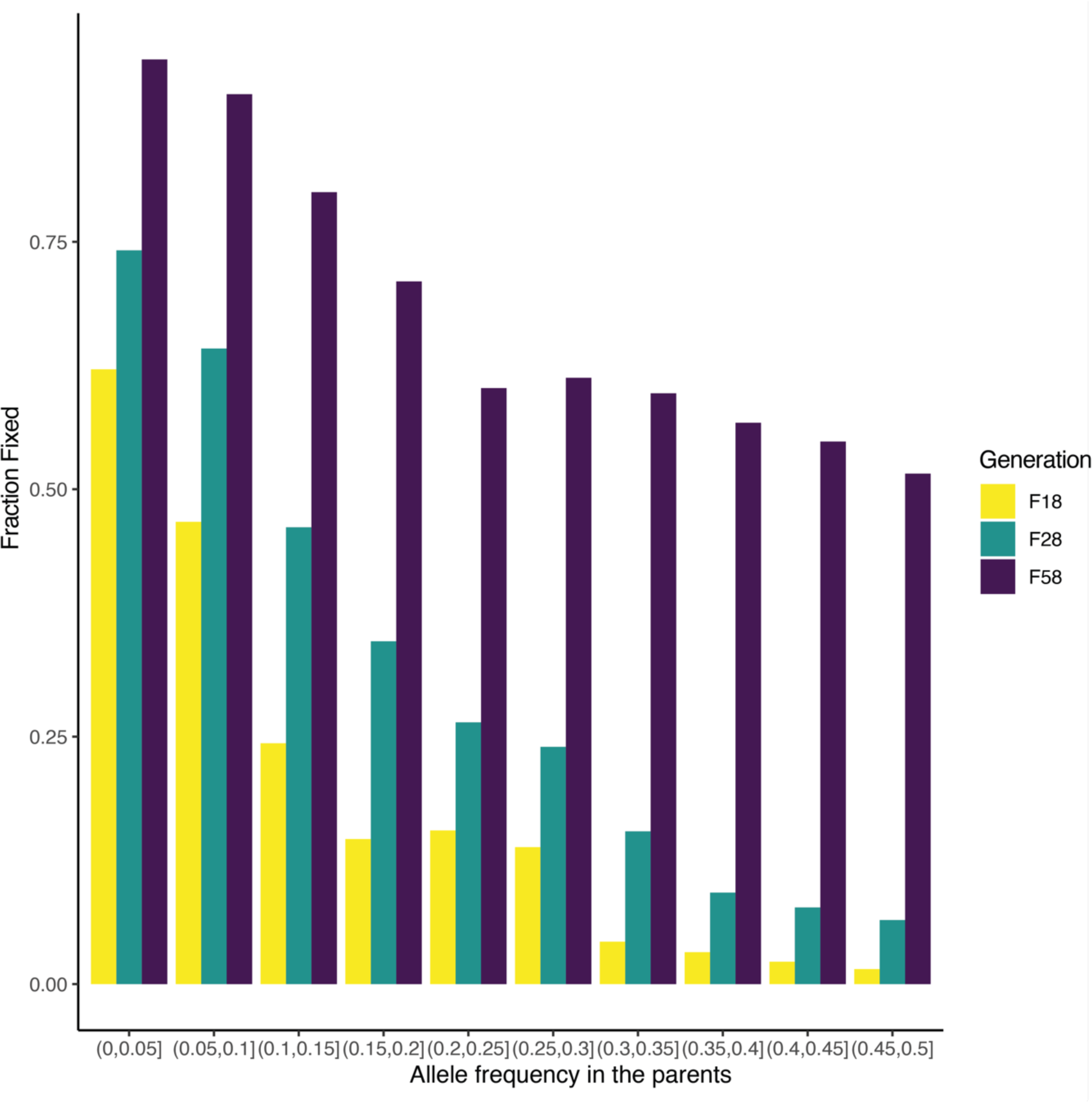
The fraction of alleles fixed over time based on their starting allele frequency. Fixation in this analysis is based on a failure to detect the allele in our sample, but the variant may occur at low frequencies.

**Fig. S7.**
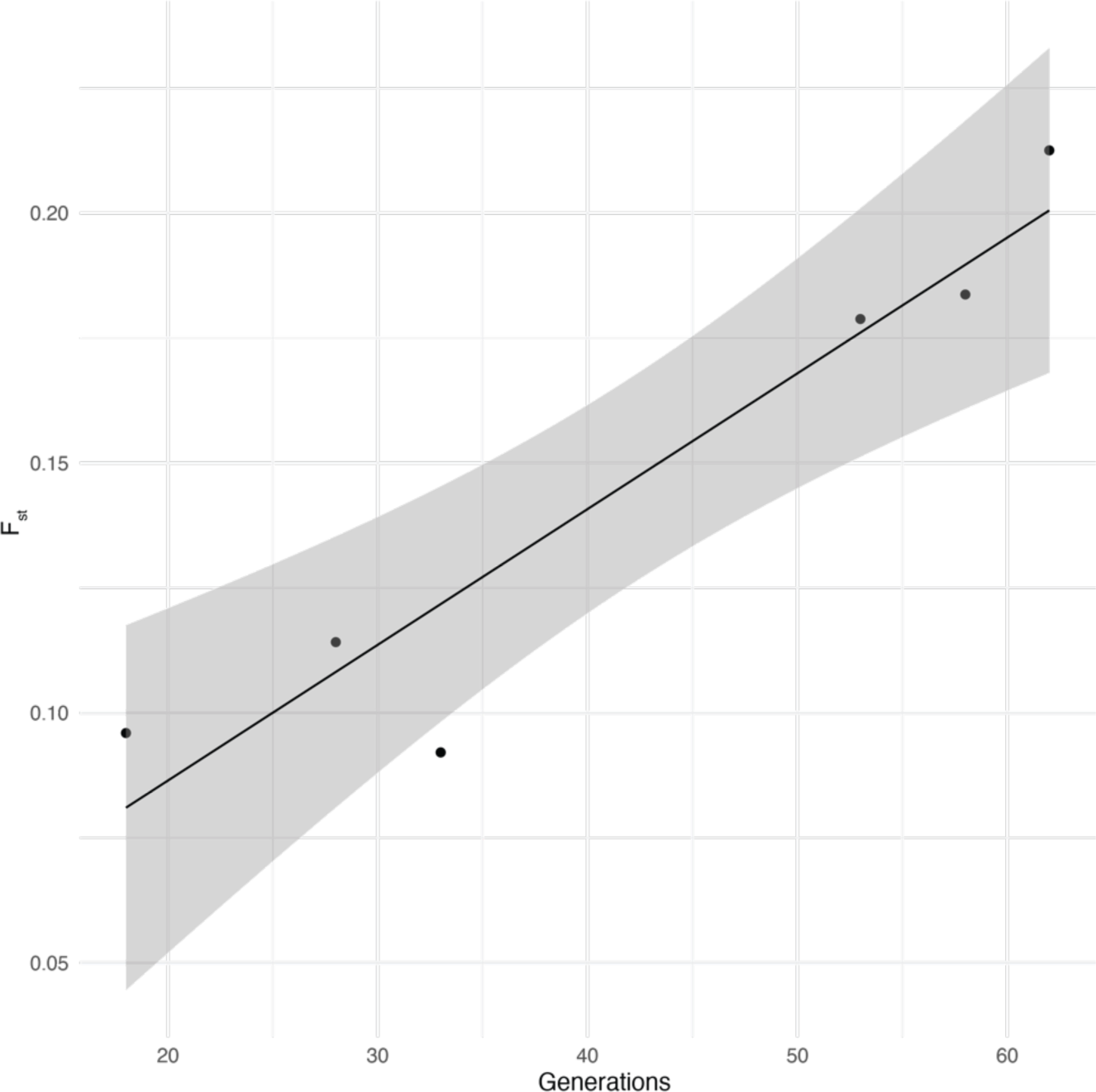
Divergence of the CCII over time. Points are the observed Fst calculated by comparing across all sampled generations and the line shows the linear regression fit.

**Fig. S8.**
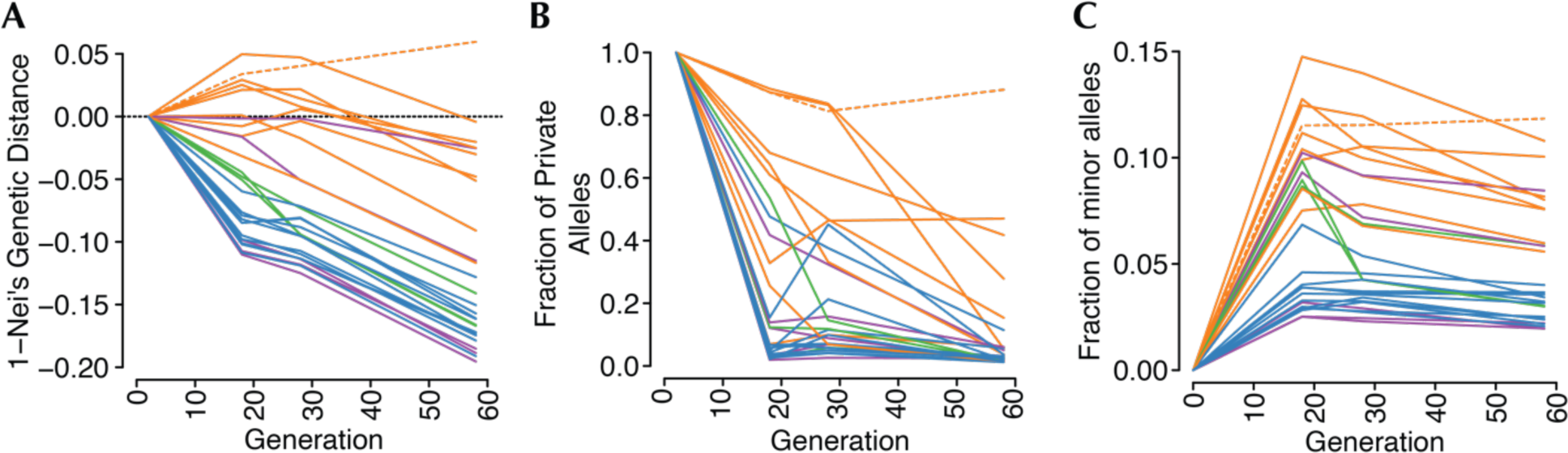
The varied success of parental alleles in the CCII. (**A**) Nei’s genetic distance calculated between each parent and the CCII at each timepoint. Each line represents a parental accession. (**B**) Fraction of private alleles specific to each parent found at allele frequency > 0 over time. (**C**) The fraction of minor alleles carried by a parent that increased in frequency relative to the founders.

**Fig. S9.**
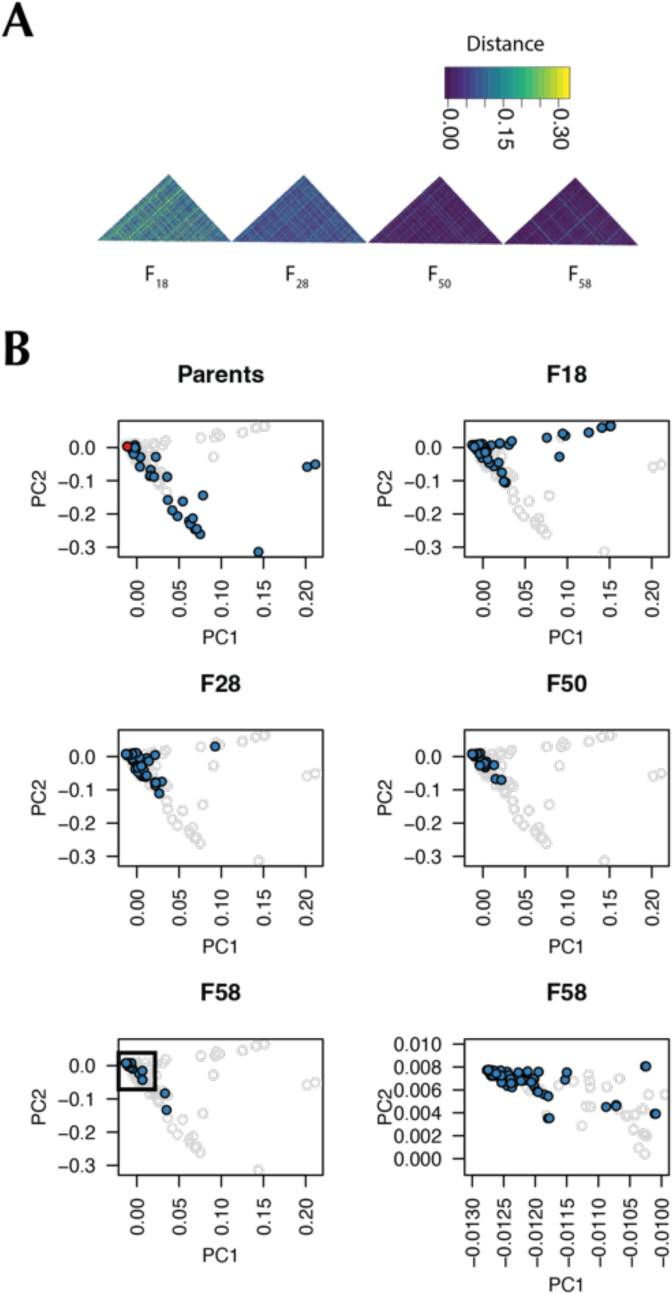
Genetic diversity in the CCII over time calculated from individual genotypes. (**A**) Heatmap showing the progressive reduction in hamming distance between progeny over time. (**C**) Principal component analysis of the genotyped CCII parents and progeny showing. The red point in the parent’s plot is Atlas. In generation F58, nearly all individual cluster in the top left corner of the plot near Atlas (boxed region is blown up in the bottom right plot).

**Fig. S10.**
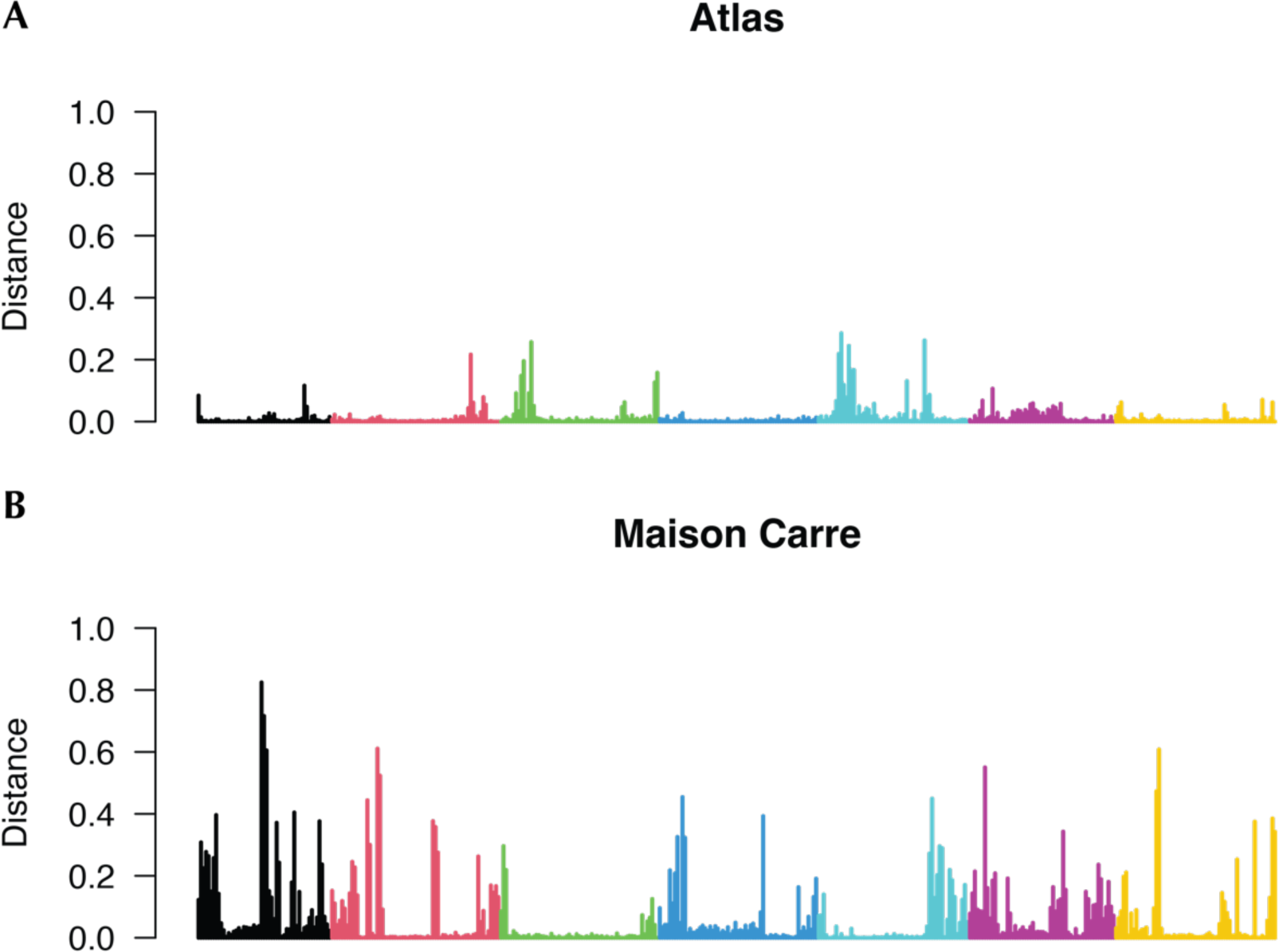
The ancestry of Lin^1^. Windowed genetic distance between the inferred primary and secondary parents of Lin^1^. Window size was 10Mb without overlaps. (**A**) Atlas shows extensive similarity to Lin^1^, but they are not identical. (**B)** Notable large regions of similarity to Maison Carree can be seen at the top of chr3H, and in the pericentromere of chr5H. Smaller segments of identity are masked with large windows sizes like those used here. The bias toward Atlas alleles observed in this dataset may reflect strong selection in the earliest generations where families segregated for alleles from both parents or subsequent outcrossing events with other progeny with Atlas ancestry.

**Fig. S11.**
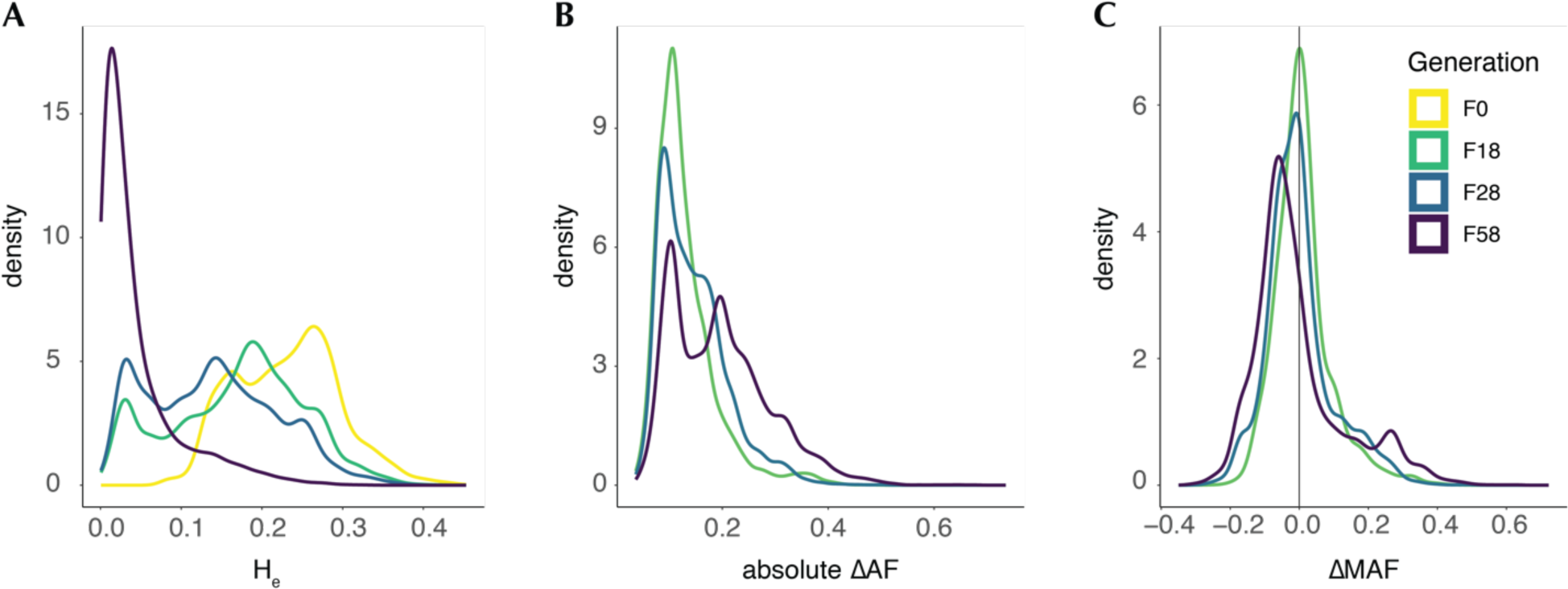
The distribution of diversity and allele frequency change statistics in genomic windows. (**A**) Windowed He (**B**) absolute change in allele frequency and (**C**) directional change in frequency of the parental minor allele.

**Fig. S12.**
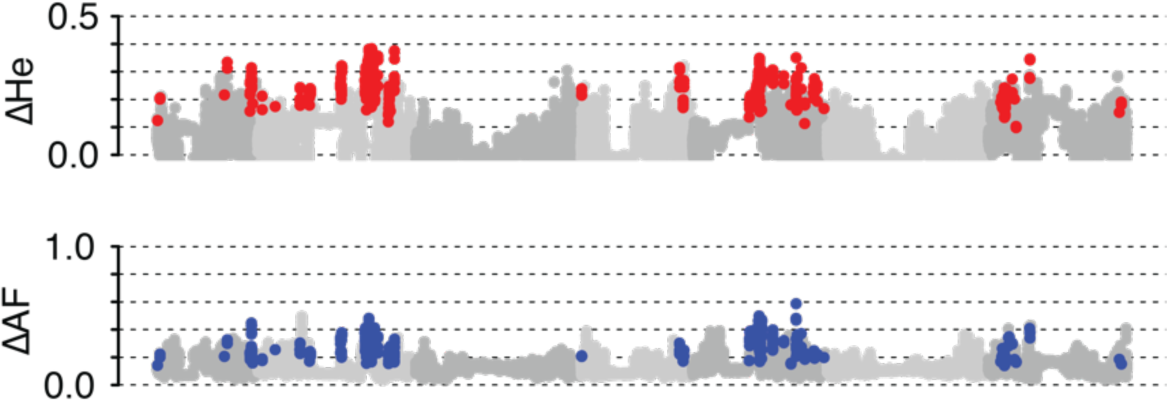
Genome wide distribution of change in He and allele frequencies from the CCII founders and F18. Colored points are windows overlapping with selected regions.

**Fig. S13.**
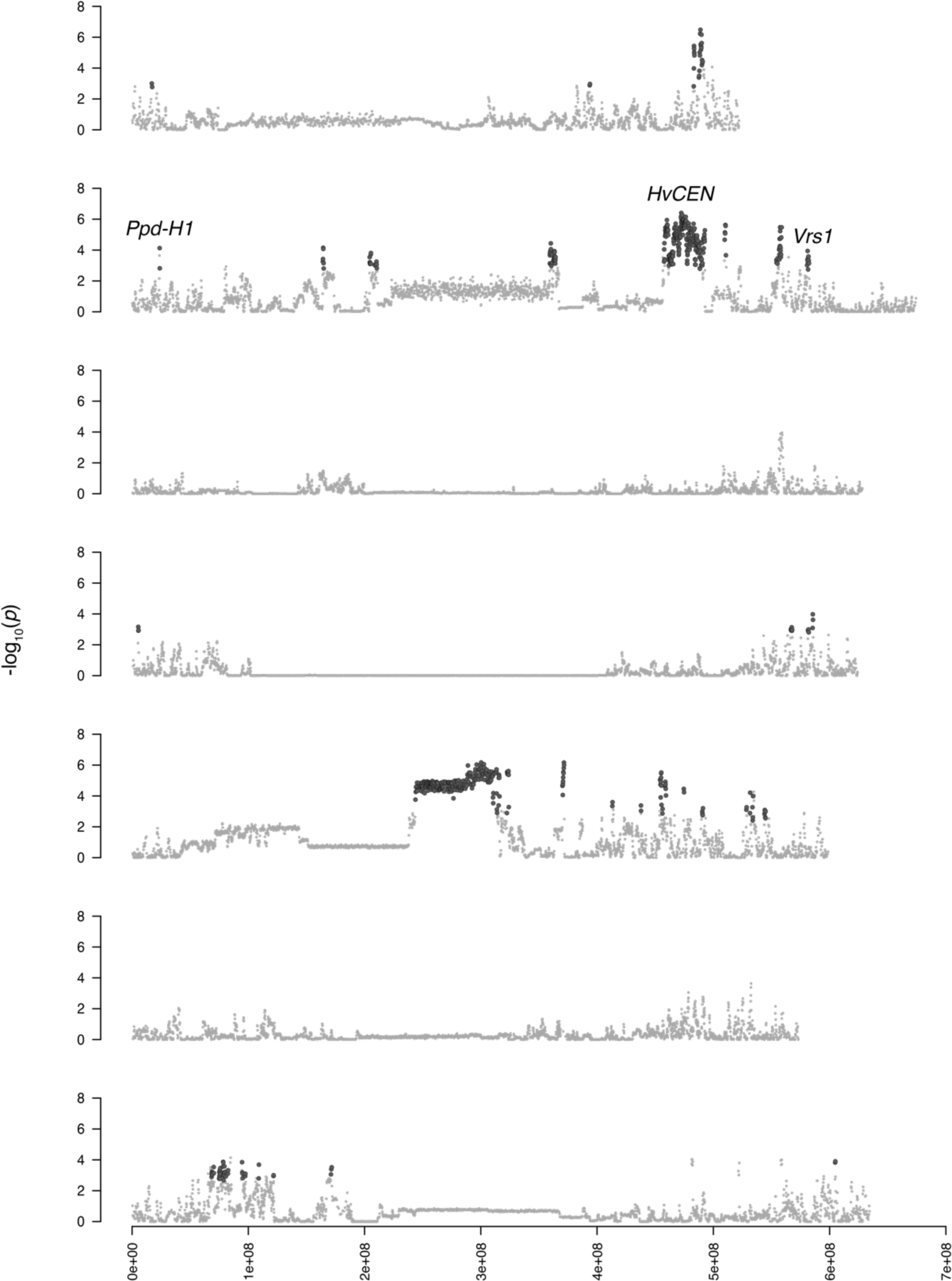
Selection scans from Fig. 3 plotted by chromosome for clarity.

**Fig. S14.**
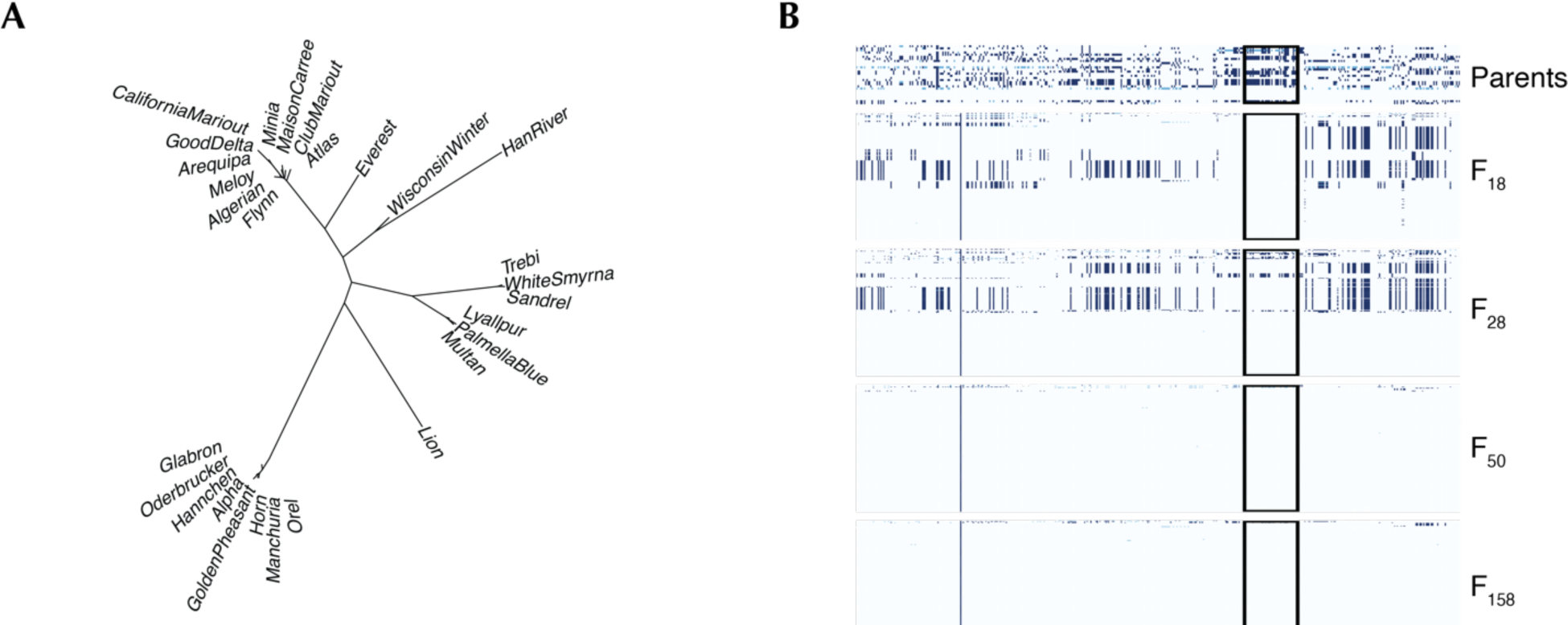
Haplotype structure around *Ppd-H1* in the CCII. (**A**) Neighbor joining tree built from parental genotypes at *Ppd-H1* including the 10kb upstream and downstream of the gene. (**B**) Haplotype structure around *Ppd-H1* across the whole CCII experiment. Box highlights the selected region.

**Fig. S15.**
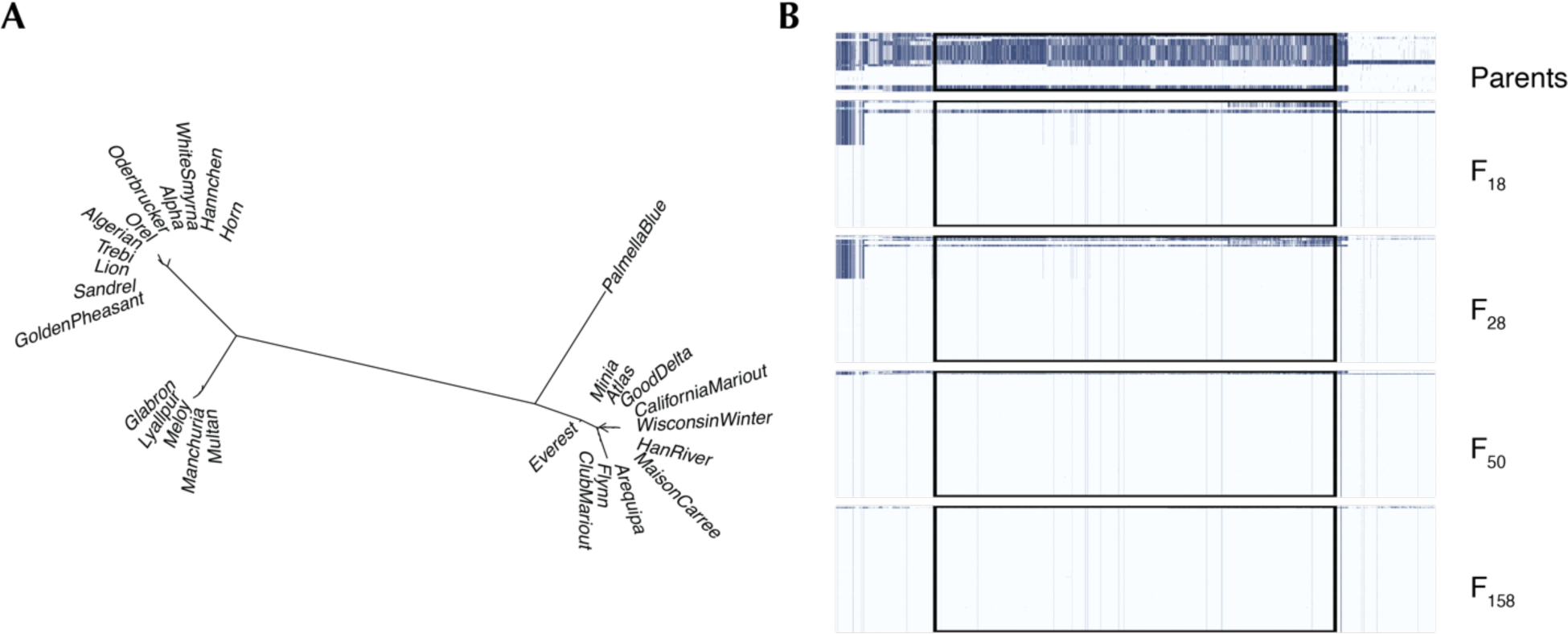
Haplotype structure around *HvCEN* in the CCII. (**A**) Neighbor joining tree from built parental genotypes at *HvCEN* including the 10kb upstream and downstream of the gene. (**B**) Haplotype structure around *HvCEN* across the whole CCII experiment. Box highlights the selected region.

**Fig. S16.**
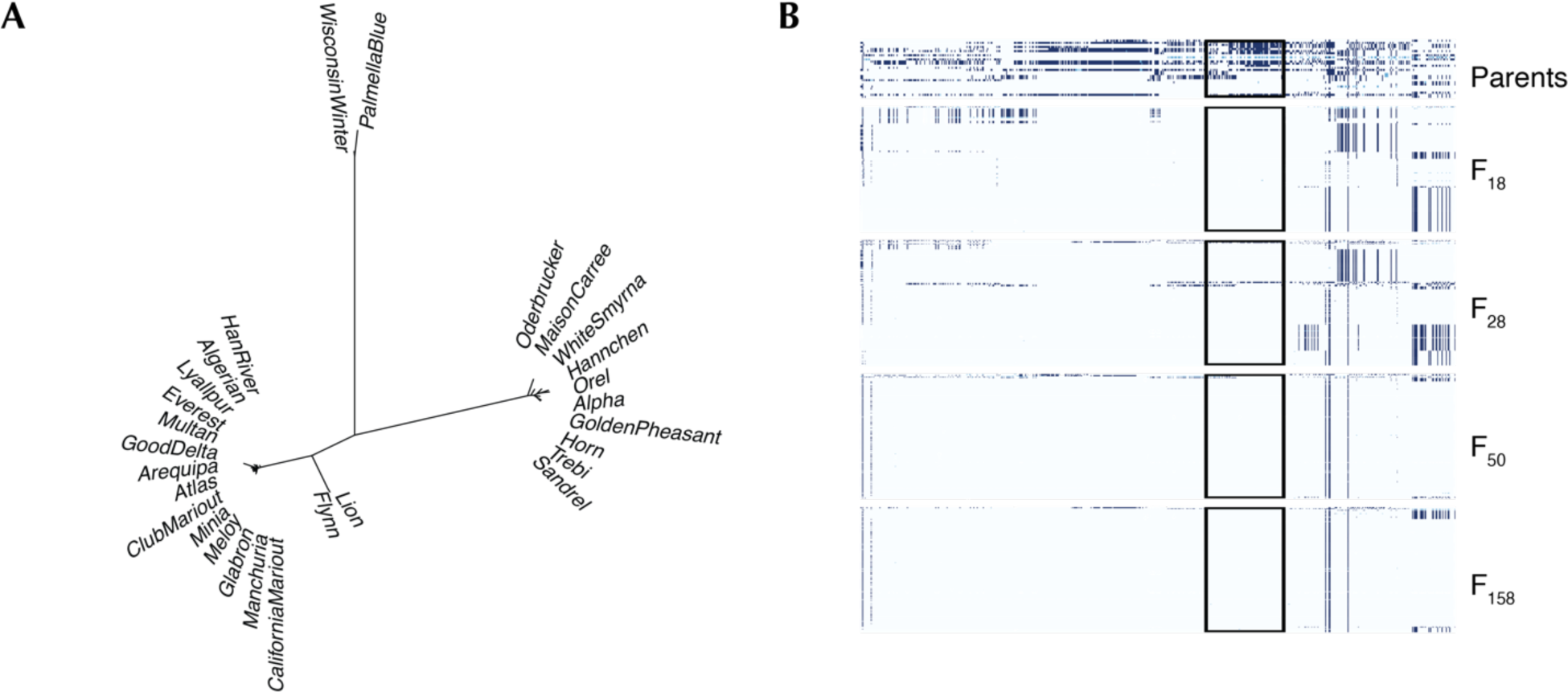
Haplotype structure around *Vrs1* in the CCII. (**A**) Neighbor joining tree built from parental genotypes at *Vrs1*including the 10kb upstream and downstream of the gene. (**B**) Haplotype structure around *Vrs1* across the whole CCII experiment. Box highlights the selected region.

**Fig. S17.**
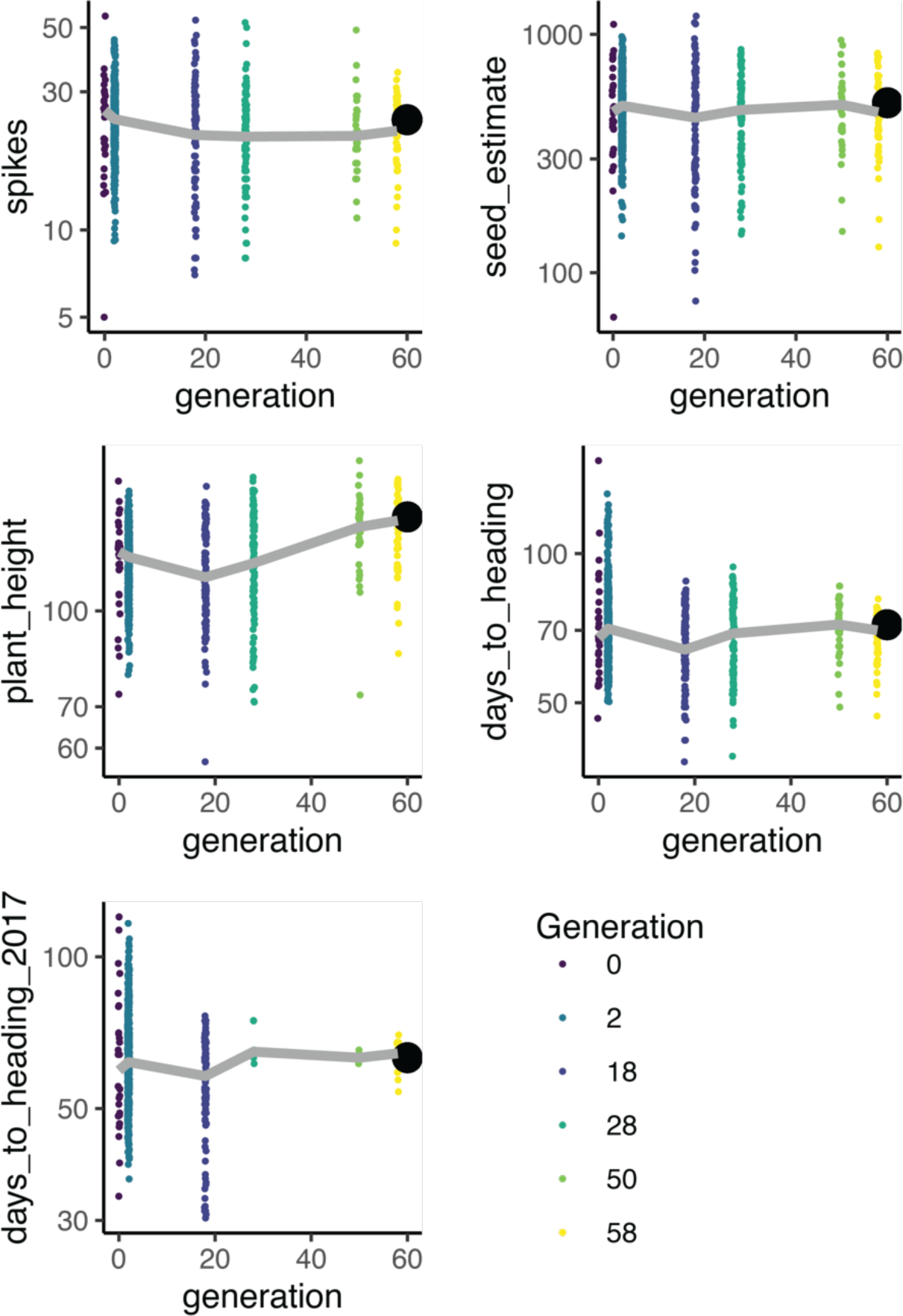
Adaptive trait shifts in the CCII. Distributions of trait values across the sampled generations in the CCII. Generation F2 values were calculated from the parental values as midparent means. All traits were measured in the greenhouses in 2016, except for the replicated measurement of days to heading which was remeasured in a subset of lines in 2017.

**Fig. S18.**
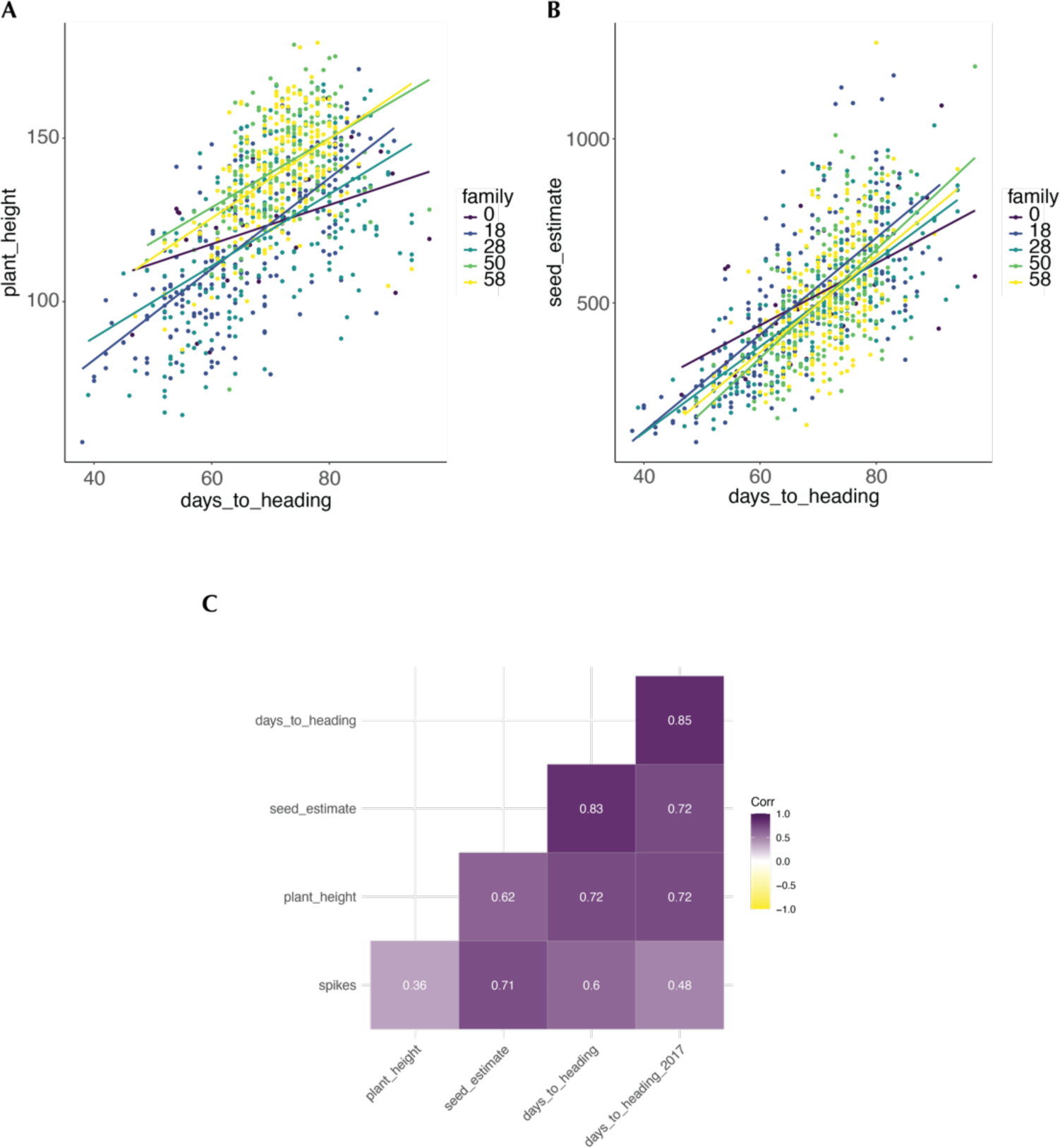
Fitness related trait correlations in the CCII. Correlation between days to heading and (**A**) plant height and (**B**) fecundity. Linear model fits are shown as solid lines. Fits were similar within each generation except for the parents which tended to show a weaker relationship. The parents still contained vernalization dependent winter varieties that were eliminated from the CCII. These varieties did not flower robustly under our greenhouse conditions. (**C**) Spearman’s correlation between all analyzed traits in this study.

**Fig. S19.**
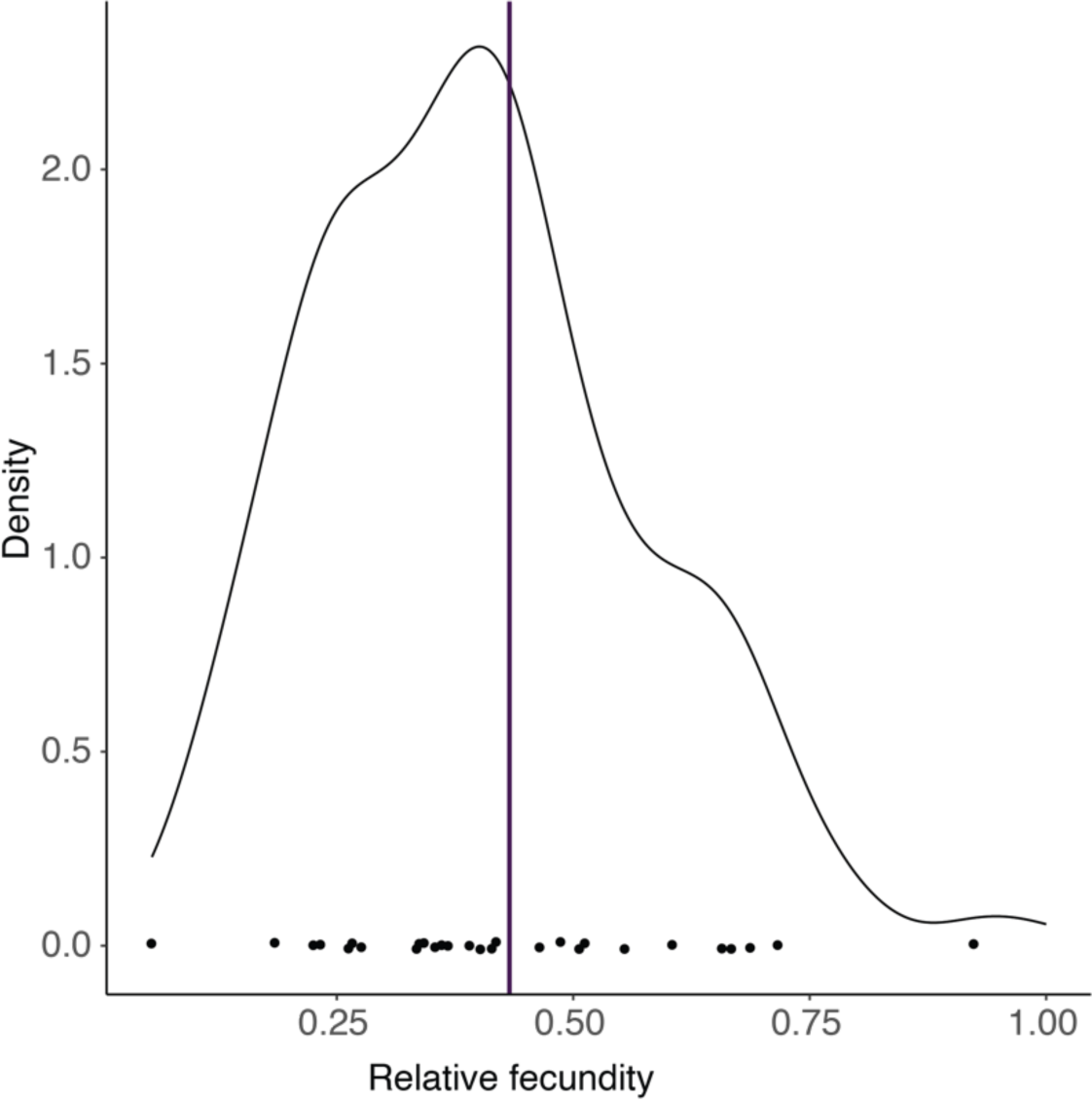
The distribution of relative fecundity of the CCII progeny in the greenhouse. Points show the mean values of the 28 parents and the line shows the value for Lin^1^.

**Fig. S20.**
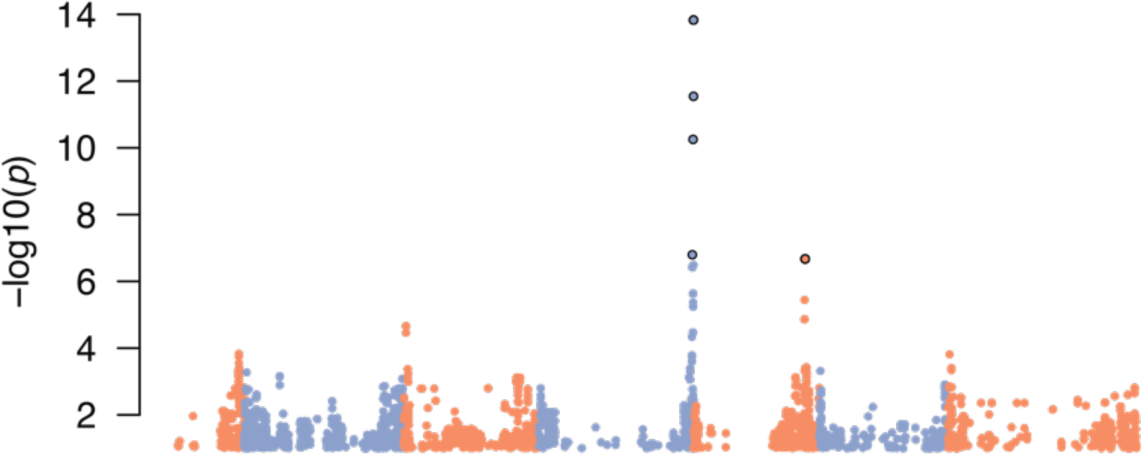
Replicated days to heading significant association on the long arm of chromosome 4H in 2017.

**Fig. S21.**
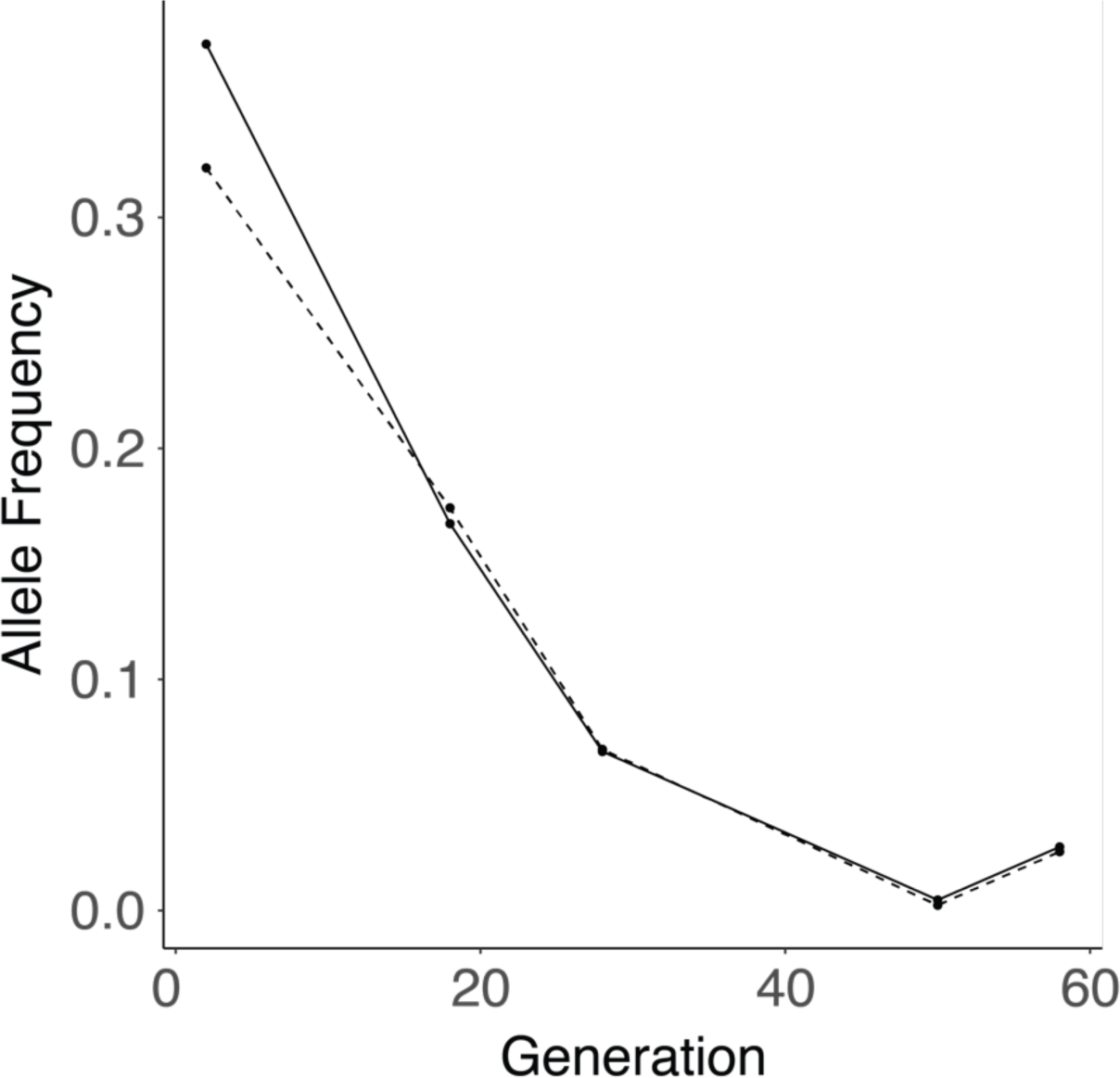
A shift in the frequency of Vrn-H2 linked SNPs over time. Solid line is the tag SNP chr4H:624173658 and the dashed line is the significant SNP chr4H:618589584 nearest to Vrn-H2.

**Fig. S22.**
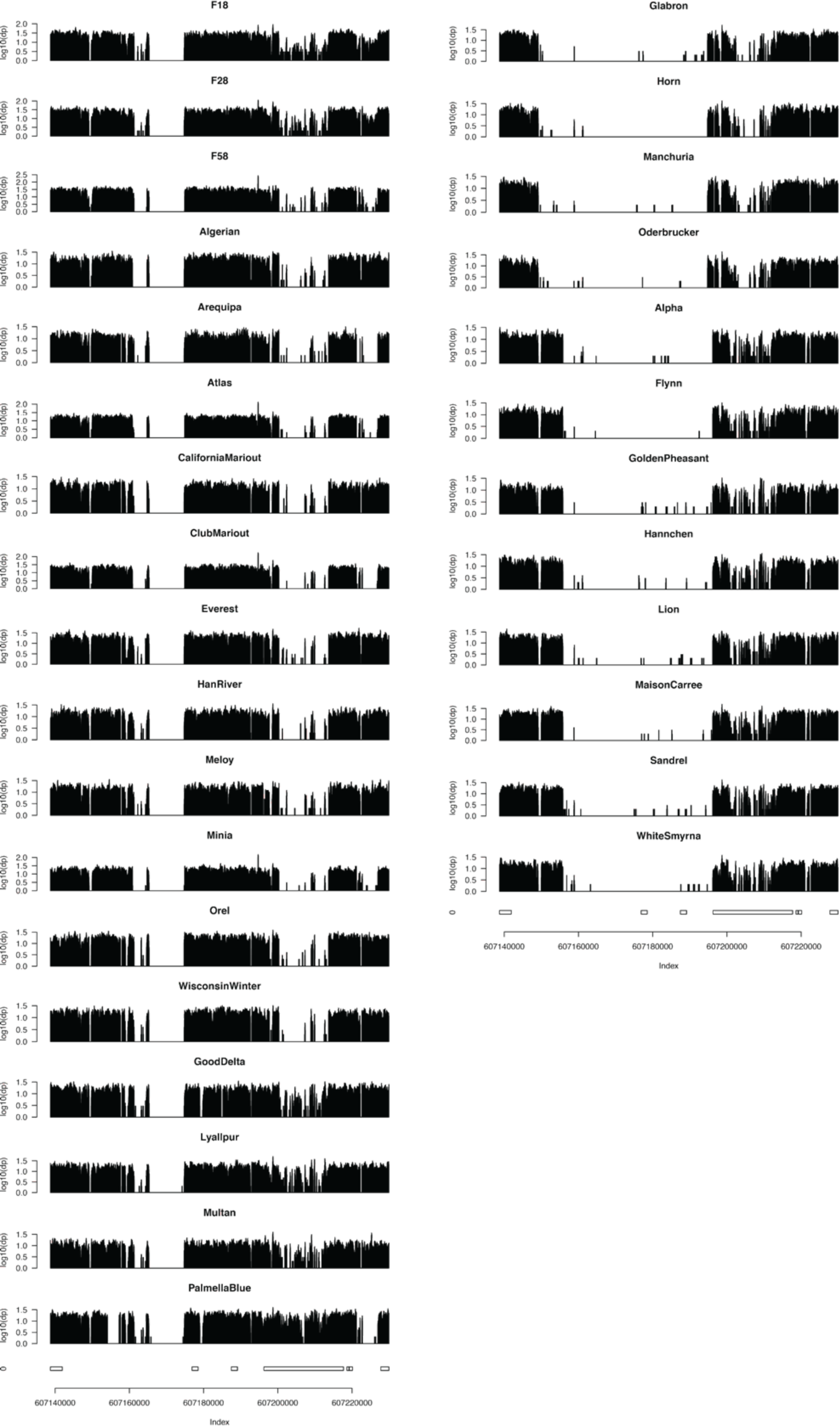
Coverage across the Vrn-H2 region for all of the CCII parents.

**Fig. S23.**
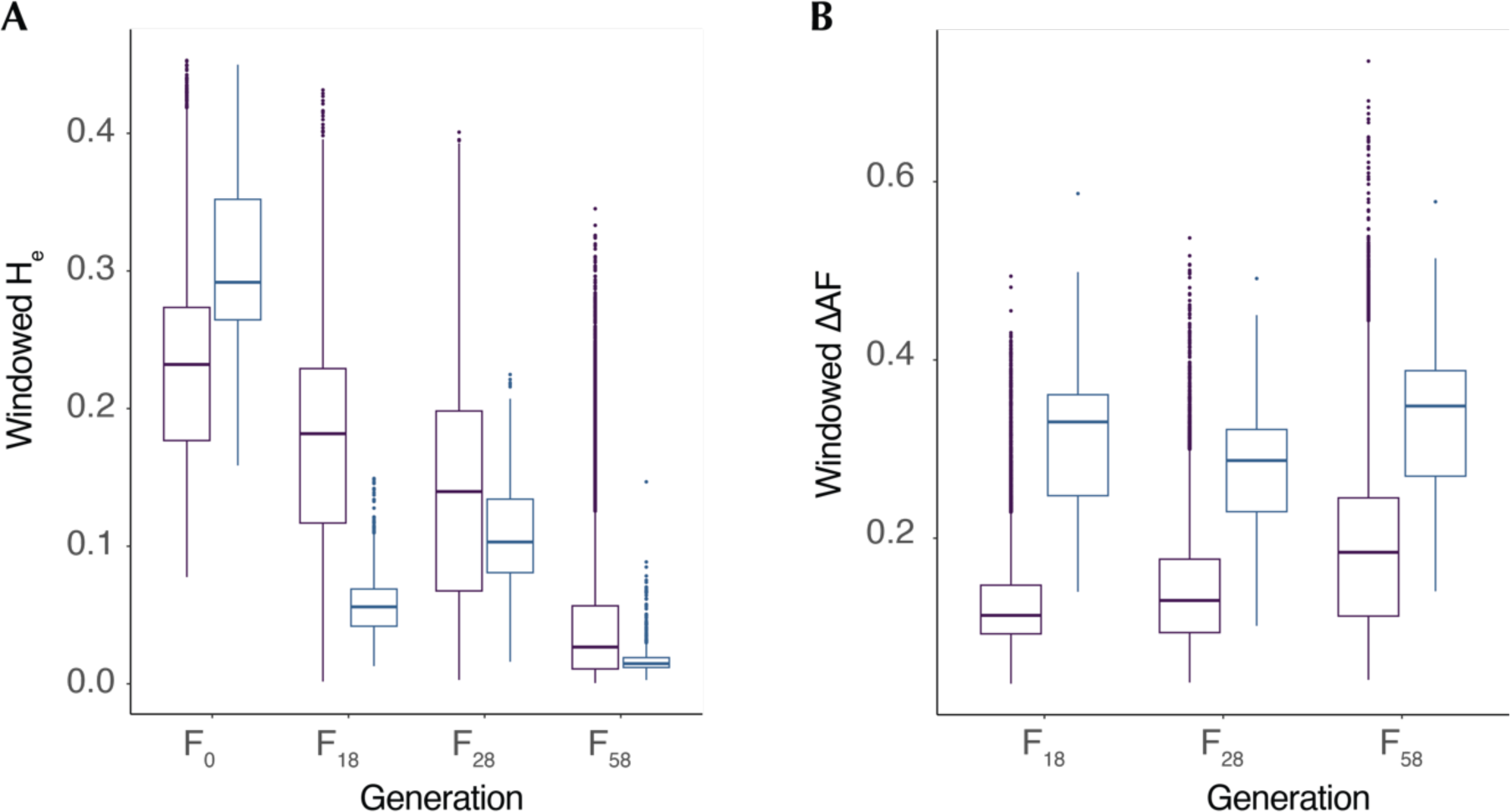
The footprint of selection over time. (**A**) The distribution of median He in genomic windows over time in genomic background (purple) and selected regions (blue). (**B**) The distribution of median absolute change in allele frequency in windows over time.

